# An alternative AUG codon in segment 5 of the 2009 pandemic influenza A virus is a swine-derived virulence motif

**DOI:** 10.1101/738427

**Authors:** Helen M. Wise, Eleanor Gaunt, Jihui Ping, Barbara Holzer, Seema Jasim, Samantha J. Lycett, Lita Murphy, Alana Livesey, Russell Brown, Nikki Smith, Sophie Morgan, Becky Clark, Katerine Kudryavtseva, Philippa M. Beard, Jonathan Nguyen-Van-Tam, Francisco J. Salguero, Elma Tchilian, Bernadette M. Dutia, Earl G. Brown, Paul Digard

**Author notes:** Equal contributions (Helen M. Wise, Eleanor Gaunt). HMW: Clinical biochemistry, Clock Tower building, Western General Hospital, Edinburgh, EH4 2XU, UK. JP: Institute of Immunology, College of Veterinary Medicine, Nanjing Agricultural University, China. SJ: MRC-University of Glasgow Centre for Virus Research, Glasgow, G61 1QH, UK. SM: Respiratory Medicine Unit, NDM Experimental Medicine, University of Oxford, John Radcliffe Hospital, Oxford, OX3 9DU, UK. Corresponding author: Paul Digard.

## Abstract

The 2009 influenza A virus (IAV) pandemic (pdm2009) was caused by a swine H1N1 virus with several atypical genetic features. Here, we investigate the origin and significance of an upstream AUG (uAUG) codon in the 5’-untranslated region of the NP gene. Phylogeny indicated that the uAUG codon arose in the classical swine IAV lineage in the mid 20^th^ Century, and has become fixed in the current triple reassortant, variant pdm2009 swine IAV and human pdm2009 lineages. Functionally, it supports leaky ribosomal initiation *in vitro* and *in vivo* to produce two isoforms of NP: canonical, and a longer “eNP”. The uAUG codon had little effect on viral gene expression or replication *in vitro*. However, in both murine and porcine models of IAV infection, removing the uAUG codon gene attenuated pdm2009 virus pathogenicity. Thus, the NP uAUG codon is a virulence factor for swine IAVs with proven zoonotic ability.

## Introduction

The 2009 influenza A virus (IAV) pandemic was caused by a swine-origin H1N1 (pdm2009) virus that, although highly transmissible, was markedly less pathogenic and caused substantially lower mortality than 20^th^ Century pandemic strains. Notwithstanding marked regional variation in the incidence of severe disease, estimates place the overall human mortality burden from the pandemic phase of pdm2009 at a similar level to disease caused by the preceding seasonal strains (1,2). Initial sequencing of the pdm2009 virus highlighted several features that could potentially explain this unexpectedly mild pathogenicity phenotype (3). These included a PB2 subunit of the viral RNA polymerase with avian-signature motifs at positions 627 and 701, a disrupted PB1-F2 gene, polymorphisms in the NS1 protein that abrogated host cell translational shut-off activity and removed a PDZ-binding domain, as well as a truncated PA-X gene (3-6). A further unusual feature of the pdm2009 genome is the presence of an upstream AUG (uAUG) codon in the 5’ untranslated region (UTR) of segment 5 (7). Segment 5 encodes the viral nucleoprotein (NP); a single strand RNA-binding protein that (along with the viral polymerase) encapsidates the single-stranded IAV genomic RNA segments into ribonucleoprotein (RNP) particles and thereby plays an essential role in supporting viral RNA synthesis (8,9). NP also contains nuclear localisation (NLS) and nuclear export signals and, in concert with the viral matrix (M1) and nuclear export protein (NEP) as well as many cellular proteins, helps direct the nuclear import of the viral genome at the start of infection and its export after genome replication (9). This functional importance is reflected in a high level of sequence conservation across IAV strains (10) and unlike the viral HA, NA and NS1 proteins, length polymorphisms of NP are very rare. However, the NP uAUG codon is in frame with the main NP open reading frame (ORF) and, if used for translation initiation, would add an extra 6 amino-acids to the N-terminus of the protein. The N-terminal 20 amino acids of NP form a flexible region not visible in crystal structures of the polypeptide (11,12) and contain the primary NLS of the polypeptide responsible for nuclear import of monomeric NP and RNPs (13-15). It was therefore reasonable to hypothesise that an alteration to this region to produce an extended NP (eNP) variant would have functional consequences for the protein that could downregulate viral pathogenicity, thus providing an explanation for the unexpectedly low levels of morbidity seen during the 2009 pandemic. Here, we describe a test of this hypothesis that shows that while eNP is produced in infection and the uAUG codon does affect *in vivo* viral pathogenicity in mice and pigs, it unexpectedly acts to increase virulence.

## Results

### The NP uAUG codon is of swine IAV origin

Segment 5 of the pdm2009 virus was acquired from the H1N1 classical swine virus lineage which in turn is a descendant of the 1918 pandemic strain, which lacked the uAUG (16,17). Examination of all available IAV segment 5 sequences on the Genbank database that reported the 5’-UTR indicated that possession of the uAUG is a minority trait, with 9104 of 33622 sequenced viruses (27%) containing it. However, within this overall population, there were clear differences between viruses from different host species (Fig 1A, Table 1), with the uAUG being extremely rare in avian isolates (∼ 0.3%) but very frequent (approaching 90%) in swine viruses. Around one third of human isolates contain the segment 5 uAUG, with the vast majority of these being pdm2009 isolates.

**Figure 1.**
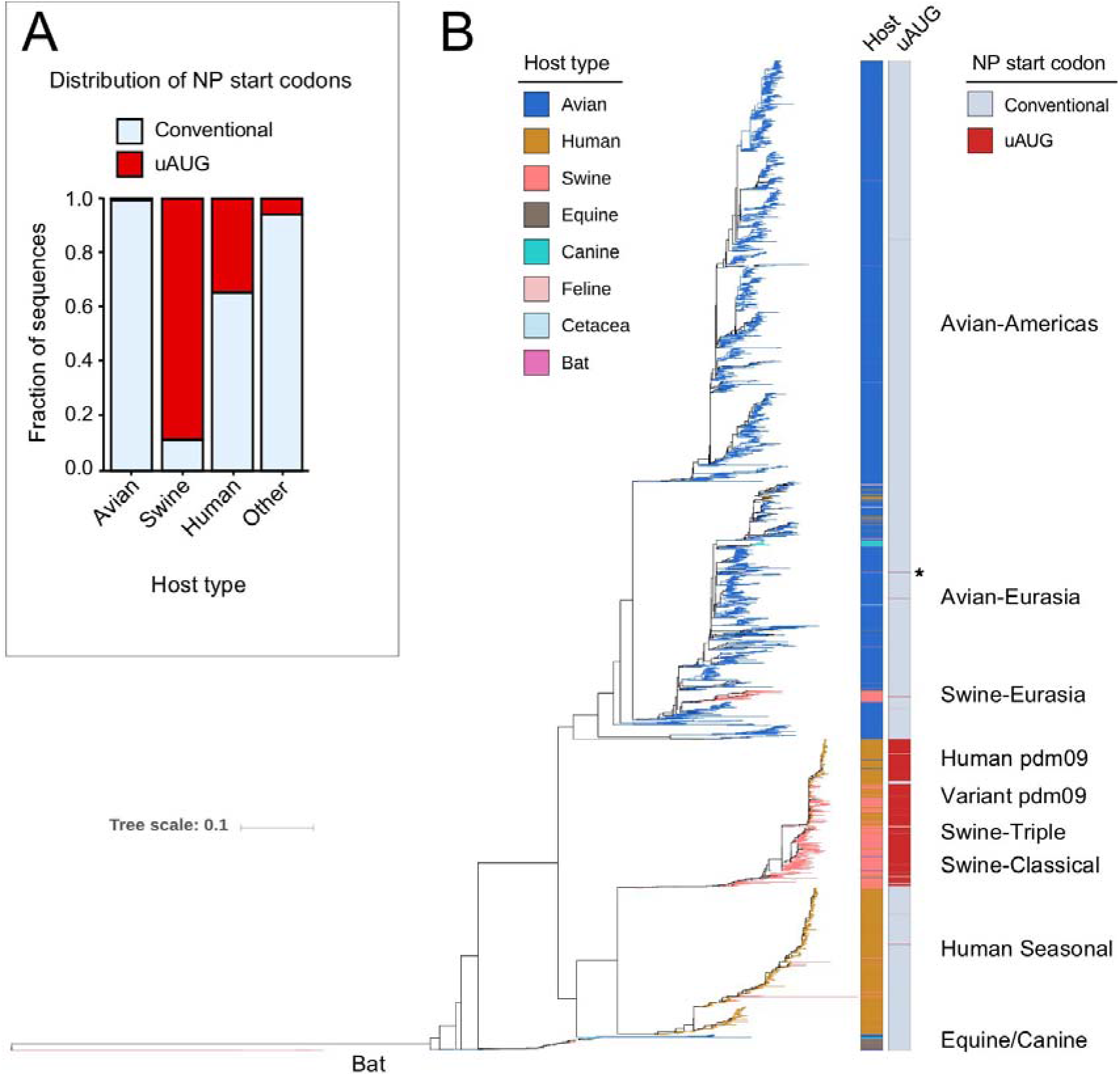
Phylogenetic analysis of segment 5 uAUG occurrence. (A) The fraction of segment 5 sequences that report the 5’-UTR, split into broad host categories, that contain the uAUG codon (red) or only the conventional NP start codon (blue). (B) Maximum likelihood phylogenetic tree of stratified subsampled sequences. Tips and left hand bar are coloured according to host while the right hand bar reports the presence (red) or absence (blue) of the uAUG start codon. Major lineages are indicated. The section of the tree indicated with an asterisk is reported in greater detail in Figure S4.

**Table 1.** Prevalence of the uAUG in IAV segment 5 sequences.

| Host | Number of sequences | Conventional Start | uAUG start (%) |
| --- | --- | --- | --- |
| Avian | 11210 | 11176 | 34 (0.3) |
| Swine | 2396 | 261 | 2135 (89.1) |
| Human | 19772 | 12852 | 6920 (35.0) |
| Other | 244 | 229 | 15 (6.1) |
| Total | 33622 | 24518 | 9104 (27.1) |

To examine the evolution of the uAUG in IAV, a phylogenetic tree for segment 5 was constructed from a stratified subsampled dataset of over 6000 sequences (Tables S1, S2) and coloured according to host species and the presence or absence of the uAUG (Fig 1B). This indicated that the uAUG is primarily a feature of the classical swine virus segment 5 and, by descent (18), the triple reassortant, pdm2009 and variant pandemic lineages (Table S1, Table S2, Fig S1). Within this clade, the uAUG codon evolved in the early 1960s in the US swine population, becoming predominant by the 1970s (Fig S2). The uAUG polymorphism also shows almost complete fixation in the subsequent triple reassortant, pdm2009 and variant swine virus lineages (93% of the sequences analysed; Fig S3, Table S1). In addition, the uAUG appears to have arisen independently on several other occasions: two or three times each in the avian and human seasonal H3N2 lineages, detectably persisting for no more than two or three years at most (Table S3), as well as twice within swine IAVs. One of the swine episodes reflects a relatively short-lived occurrence, in which an H5N1 virus transferred from ducks to pigs (19), gaining the uAUG codon around the time of the epizootic transition (Fig S4). The other occasion represents a localized gain of the uAUG within the Eurasian swine IAV lineage in Hong Kong in the early/mid 2000s (Fig S5). Thus overall, segment 5 has gained the uAUG codon on at least seven occasions; three of these were associated with swine IAV and the first of these, acquired in the background of a segment from the 1918 pandemic virus, has persisted for over half a century and resulted in world-wide colonization of swine, and via the 2009 pandemic, man.

**Table 2.**
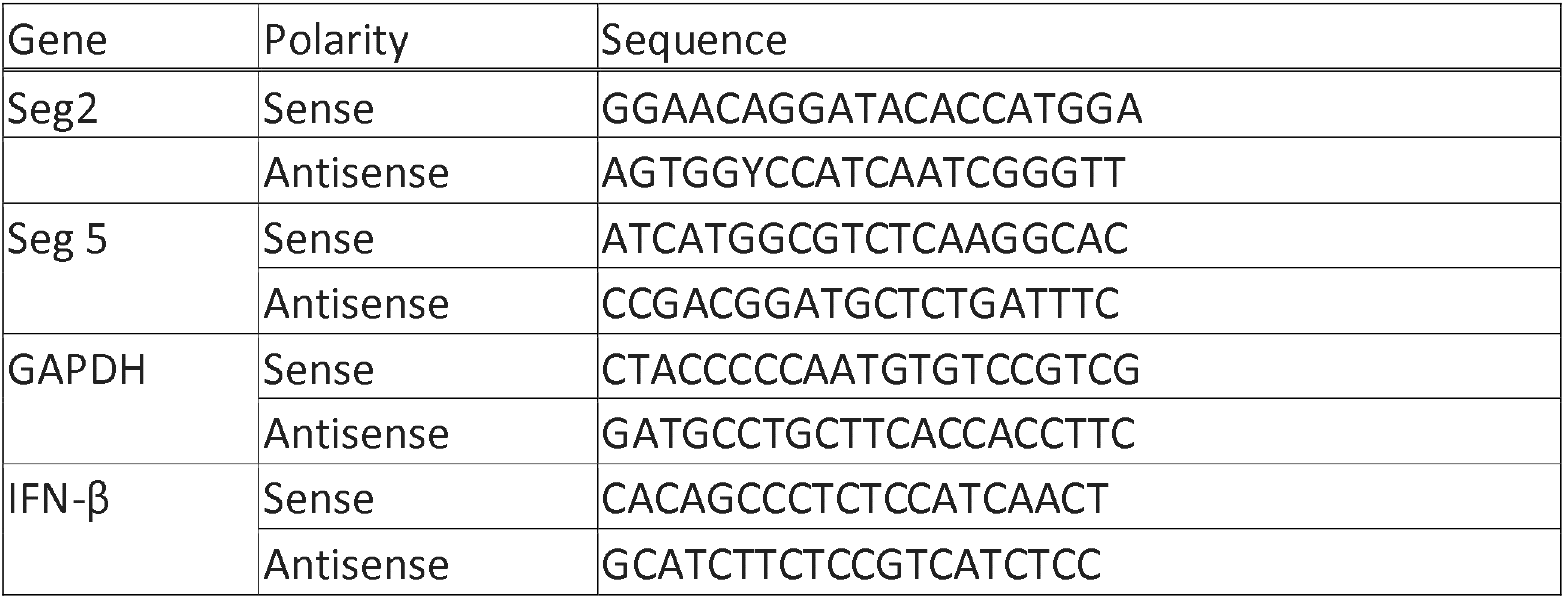
Sequences of primers used for RT-qPCR.

### Initiation of translation occurs from the segment 5 uAUG in cell-free and cell-based systems

The phylogenetic data suggested the hypothesis that the uAUG provided a host-specific selective advantage in H1N1 viruses. As a first test of its biological significance, we asked if it was used for translation. The uAUG arises from a C28A polymorphism and is in frame with the canonical start codon of the NP ORF (AUG1) such that, if used, it would produce an extended NP polypeptide with a 6 amino-acid extension (Figs 2A, B). However, the uAUG codon is in a poor ‘Kozak’ context for translation initiation (20) so it was unclear if, or to what extent, it might be seen by scanning ribosomes and used for translation initiation. A similarly poor context AUG codon near the 5’-end of segment 2 of IAV is not seen by scanning ribosomes to any appreciable extent (21). To address whether the segment 5 uAUG is used for translation, we created a series of constructs based on segment 5 cDNA, from either the UK prototype pdm2009 virus A/England/195/2009 (Eng195) (22) or the laboratory-adapted A/Puerto Rico/8/34 (PR8) H1N1 strain, with mutations designed to alter potential translation start sites in the first 100 nucleotides. These included removing the uAUG from Eng195 or adding it to PR8 by A28C/C28A switches, altering AUGs 1 and 2 to CUG codons, improving the Kozak consensus sequence of uAUG (by U25G) and reciprocally swapping the context of AUG1 (by mutating nucleotides 40, 43 and 44) between Eng195 and PR8 identities (Fig 2A).

**Figure 2.**
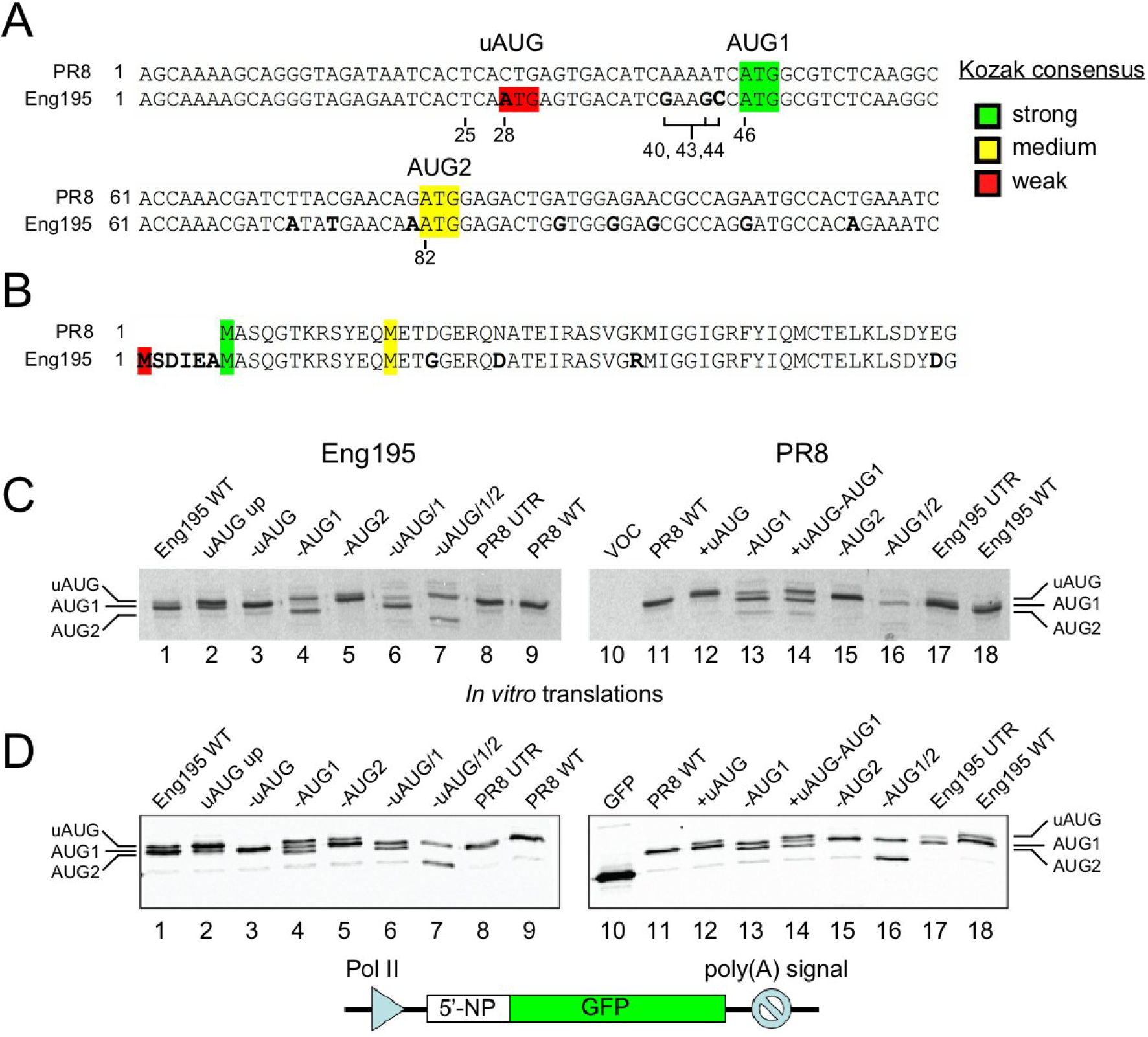
Sequence and translation initiation potential of IAV segment 5. (A) Nucleotide sequence of the 5’-end (mRNA sense, starting from position 1 of cRNA; *i.e.* ignoring any cap-snatched leader sequence) of segment 5 from PR8 and Eng195 strains. Differences are highlighted in bold, while AUG codons are colour-coded according to their Kozak initiation potential. Numbered positions were targeted for mutagenesis. (B) Amino-acid sequence of the N-terminal region of NP, highlighted as above. (C) Aliquots of rabbit reticulocyte lysate coupled *in vitro* transcription/translation reactions supplemented with ^35^S-methionine were programmed with pDUAL plasmids containing cDNA copies of the indicated WT and mutant segment 5s (or empty vector; VOC) before separation by SDS-PAGE. Radiolabelled translation products were detected by autoradiography. (D) Lysates from 293T cells transfected with plasmids containing the 5’-201 nucleotides of segment 5 cDNA, either WT or mutated as labelled, fused in frame with GFP (or with a plasmid only encoding GFP) were separated by SDS-PAGE and western blotted for NP. The migration position of polypeptides starting at the first three AUG codons is indicated.

These plasmids were then used to programme radiolabelled coupled *in vitro* transcription-translation reactions in rabbit reticulocyte lysate and polypeptide synthesis was monitored by SDS-PAGE (run for longer than normal to separate polypeptides predicted to differ in molecular weight by < 1 kDa) and autoradiography. The wild type (WT) PR8 plasmid directed synthesis of a single major polypeptide species whereas WT Eng195 gave a doublet in which the slower migrating species was less abundant (Fig 2C, compare lane 1 with 9, and lane 11 with 18), consistent with translation initiation at either of two closely spaced AUG codons in the Eng195 segment 5 mRNA. Further supporting this conclusion, improving the Kozak consensus of the Eng195 uAUG altered the proportions of the doublet so that the upper band was predominant, while mutating the uAUG removed it (Fig 2C, lanes 2 and 3). Confirming the likely identity of the polypeptides, mutation of AUG1 to CUG further changed the ratio of the doublet species, with only a trace of the smaller polypeptide now visible, but with the addition of a more prominent faster migrating band (lane 4) that co-migrated with trace species visible in WT and other AUG1-containing translation reactions. Mutation of AUG2 removed this fast-migrating product (lane 5), suggesting that leaky ribosomal scanning past an inefficiently recognised uAUG in the absence of AUG1 led to increased usage of AUG2 for translation initiation. The small amount of a polypeptide with the expected size for canonical NP seen when AUG1 was replaced with CUG most likely arose from non-AUG initiation at a CUG codon in a strong Kozak consensus (23,24), as the alternative mutation of AUG -> AGG blocked its formation (data not shown). Pairwise knockouts of the first three AUG codons in Eng/195 segment 5 also indicated leaky ribosomal scanning leading to context-dependent recognition of all three start sites (lanes 6 and 7). Creation of the equivalent mutations in the PR8 NP gene showed that similar rules applied; addition of uAUG led to production of a closely spaced NP doublet (Fig 2C, compare lanes 11 and 12), while mutating combinations of uAUG, AUG1 and AUG2 showed that the hierarchy of translation initiation potential *in vitro* was AUG1 > uAUG >> AUG2 (lanes 13-16). However, swapping the entire 5’-UTRs of Eng/195 and PR8 had little effect beyond that of the addition or omission of the uAUG (compare lanes 8 and 9, 17 and 18), suggesting that the nucleotide polymorphisms at positions 40, 43 and 44 were of little significance for initiation at AUG1.

The coupled *in vitro* transcription-translation system we used did not generate mRNAs with 5’-cap structures and nor was it optimised for KCl concentration, leading to the possibility of less accurate translation initiation than would occur in intact cells (25). We therefore tested NP expression after transfection of 293T cells with a corresponding set of plasmid constructs containing the first 201 nucleotides of segment 5 cDNA fused in frame to the green fluorescent protein (GFP) ORF (Fig 2D). This cloning strategy kept the 5’-end of the viral sequences intact but decreased the overall size of the expected polypeptides from ∼ 56 kDa to ∼ 34 kDa, thus aiding separation of the various isoforms, as well as permitting their detection by western blotting for GFP. In this system, all plasmids with an IAV UTR produced a low abundance product with the same mobility as GFP (compare lanes 10 and 11). However, in addition to this, a construct with the WT PR8 UTR produced a single major species while the WT Eng195 plasmid produced a clearly separated doublet (Fig 2D, compare lanes 1 and 8, and lanes 11 and 18). As before, the relative abundance of the Eng195 upper doublet species was increased by a mutation that improved the Kozak consensus of the uAUG while its synthesis was blocked by removal of the uAUG (lanes 2 and 3). Again, similarly to the outcome of the *in vitro* translation experiments, mutation of AUG1 to CUG produced a triplet species whose upper and lower constituents could be explained by initiation at the uAUG and AUG2 in the absence of AUG1, as well as lower levels of CUG codon-directed translation initiation at the mutated AUG1 codon (compare lane 4 with lanes 5-7). Analysis of the counterpart mutations in a PR8 background produced corresponding results; introduction of the uAUG gave a doublet NP species (lane 12) while AUG2 was only used for translation initiation after mutation of AUG1 to CUG had downregulated but not abolished initiation at the canonical NP start site (lanes 13-16). Thus, the NP uAUG codon was seen by scanning ribosomes in a cellular context as well as *in vitro* to produce eNP, although AUG1 remained the preferred start site. In contrast to the cell-free setting, translation initiation at AUG2 could only be detected in the absence of AUG1.

### eNP is functionally equivalent to canonical NP in supporting viral gene expression and replication in cells

Next, we examined the effect of a subset of these mutations on the ability of NP to support viral gene expression, using an assay in which RNPs were reconstituted by transfection of cells with plasmids encoding the three subunits of the viral RNA polymerase (3P) and WT or mutant copies of the NP gene (26,27), along with a vRNA-like reporter segment with an antisense luciferase gene. First, NP expression was examined by western blotting, where again, the presence of an uAUG codon in both PR8 and Eng195 backgrounds led to production of an NP doublet whose relative abundance varied according to the strength of the uAUG Kozak consensus (Fig 3A). Reconstitution of both WT PR8 and Eng195 RNPs led to around 300-fold increases in luciferase expression compared to a control reaction lacking NP (Fig 3B). However, at a fixed dose of NP plasmid, all of the mutants had comparable activity to their WT counterpart, with no more than 2-fold differences evident. To provide a more sensitive examination of NP activity, we titrated the amounts of NP plasmid and fitted the luciferase expression values to a variable slope dose-response enzyme kinetic model. The resulting curves for both the PR8 and Eng195 sets of plasmids were very similar (Fig 3C, D) and the estimated concentrations of plasmid required for half-maximal activity were not significantly different. Thus, the precise identity of the N-terminus of NP had little influence on viral gene expression, even at limiting amounts of the protein.

**Figure 3.**
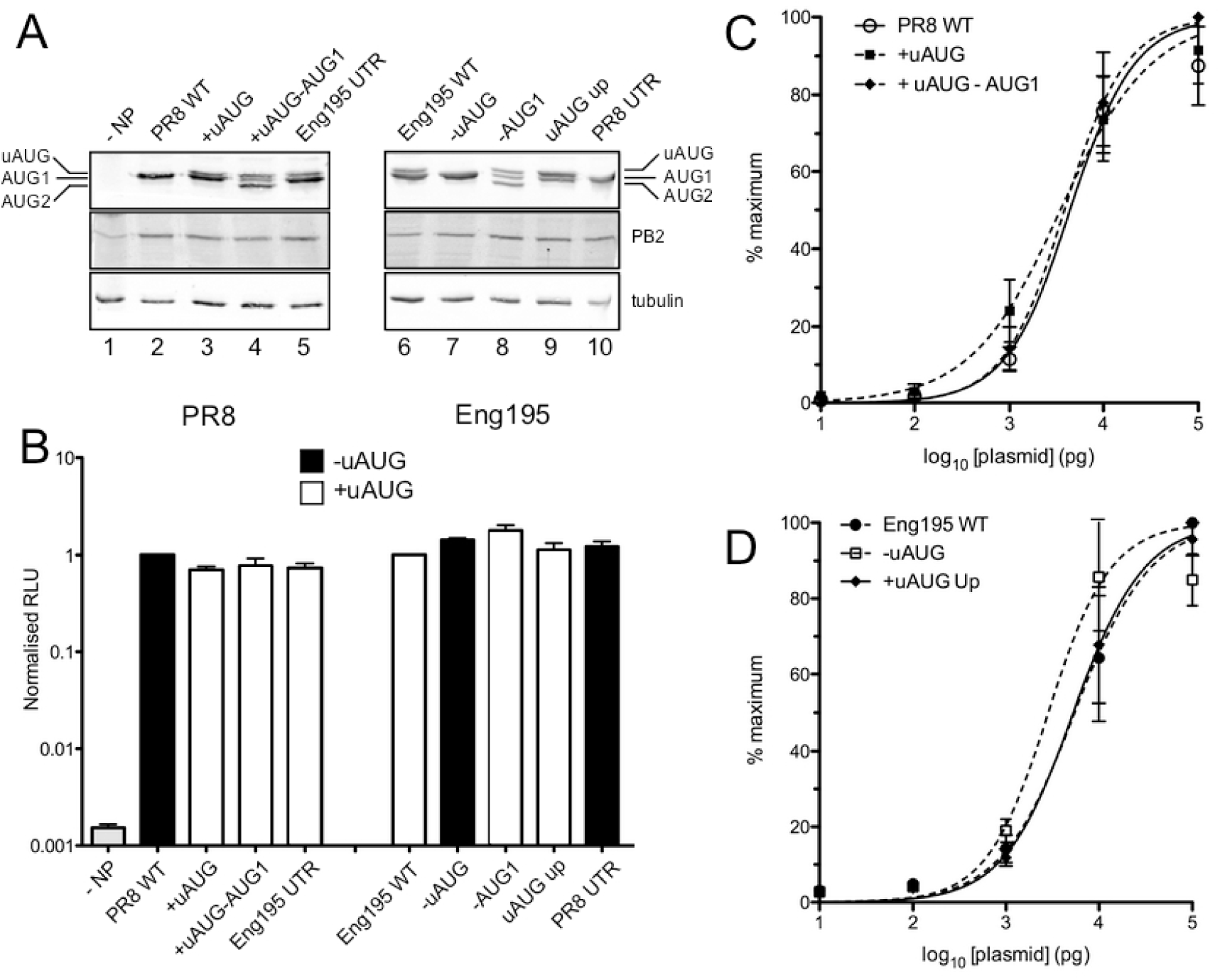
Ability of NP mutants to support viral gene expression in RNP reconstitution assays. 293T cells were transfected with reverse genetics plasmids encoding the 3 polymerase proteins (from PR8 or Eng195 as indicated), WT or mutant forms of NP and a vRNA-like reporter segment encoding luciferase, thus reconstituting RNPs. (A) Cell lysates were analysed by SDS-PAGE and western blotting for the indicated polypeptides. (B) Luciferase activity in the lysates was measured and normalised to the amount seen with the corresponding WT gene. Data are the mean ± SEM of four independent experiments. Differences between samples with a complete RNP were non-significant (repeated measures ANOVA with Dunnett’s multiple comparison test comparing against the matched WT) (C, D) Luciferase activity was measured from RNP reconstitution assays in which the NP plasmid was titrated and all other plasmids kept constant and normalised to the maximum activity within an individual titration set. Data are the mean ± SEM of 3 (PR8 + uAUG − AUG1), 4 (all Eng195 data), 5 (PR8 + uAUG) or 6 (WT PR8) independent experiments, curve fitted to a variable slope log10 [agonist]- response model using Graphpad Prism. The 95% confidence limits of the estimated EC_50_ values within groups overlapped, indicating non-significance (Table S4).

We then examined what effect the presence of the uAUG codon had on virus replication *in vitro*. End-point titres following low multiplicity infection of MDCK cells with WT PR8 or variants with the uAUG codon added to PR8 segment 5 were essentially the same (Fig 4A, left hand bars). When the counterpart experiment was performed for viruses with segment 5 from either Eng195 or another early isolate from the 2009 pandemic, A/Halifax/210/2009 [SW210; (28)] (both as 7:1 reassortants on the PR8 background to confer efficient infection of MDCK cells), removal of the uAUG codon with an A28C mutation gave slight increases (4-5 fold) in average titres while replacing the normal Eng195 UTR with the PR8 sequence gave an 8-fold increase (Fig 4A, middle and right hand bars). However, none of these differences were statistically significant. Western blot analysis of lysates from cells infected at high multiplicity confirmed that the A28C polymorphism behaved as expected with respect to production or not of the two NP isoforms in all virus backgrounds, without affecting synthesis of other viral structural proteins (Fig 4B). Since our clone of the Eng195 virus does not infect or replicate well in the continuous cell lines commonly used to study IAV replication, we tested the full SW210 virus under multicycle growth conditions in human A549 cells or swine new-born pig tracheal (NPTr) cells. WT and A28C mutant viruses replicated with almost identical kinetics in both cell types (Figs 4C, D). Thus the presence or absence of the segment 5 uAUG codon and expression of eNP had little effect on IAV replication *in vitro*.

**Figure 4.**
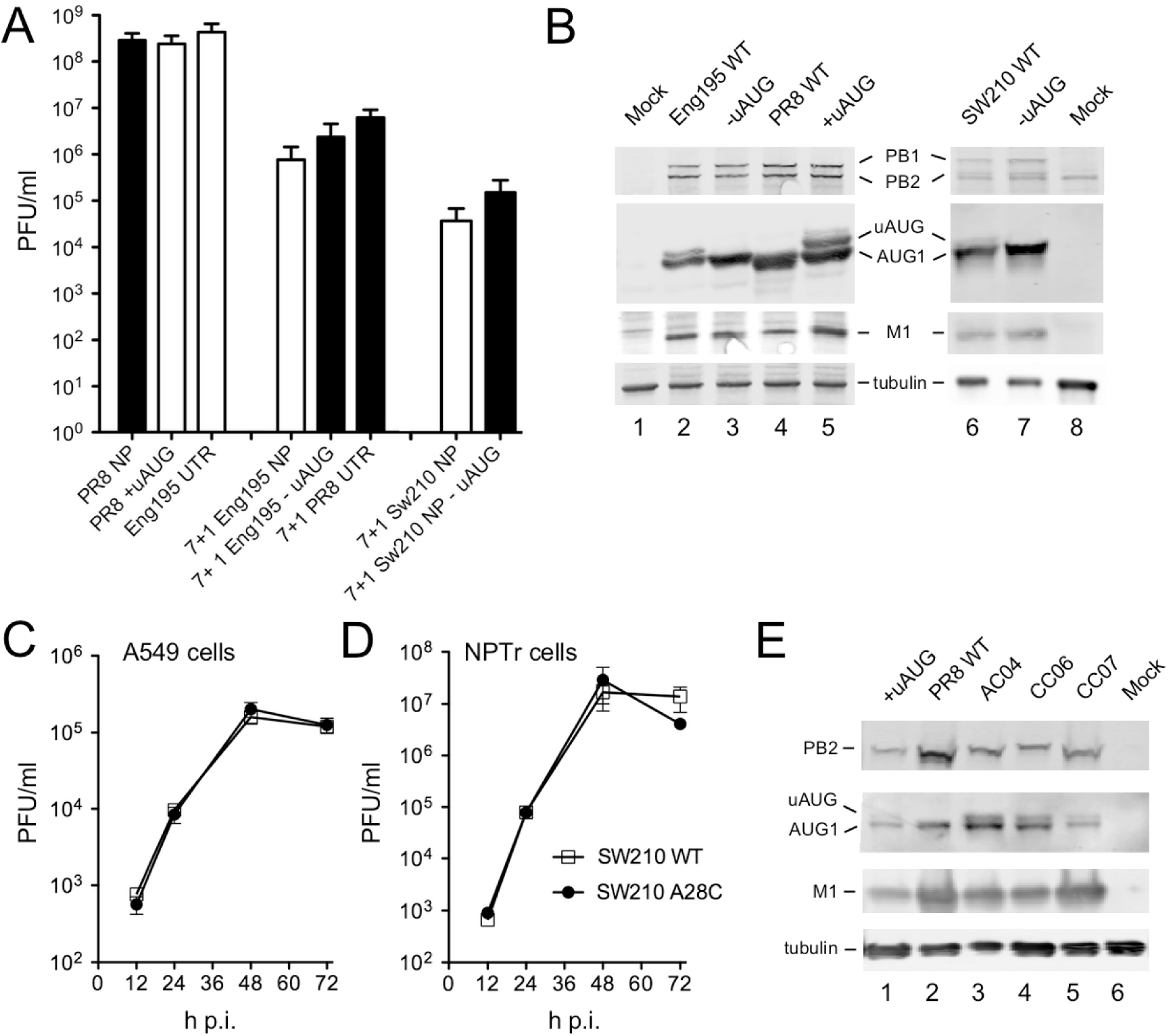
Expression and functional significance of eNP for virus replication *in vitro*. (A) MDCK cells were infected at an MOI of 0.01 with the indicated viruses and titres measured at 48 h p.i.. Data are the mean ± SEM of 4-5 independent experiments. Differences within groups were not statistically significant (PR8, Eng195; One way ANOVA with Tukey’s post test, Sw210; *t*-test). (B) Cell lysates from A549 cells infected at an MOI of 5 and harvested at 24 h p.i. were analysed by SDS-PAGE and western blotting for the indicated polypeptides (uAUG/AUG1 = NP). (C, D) A549 or NPTr cells were infected an an MOI of 0.03 and samples titred at the indicated times p.i. Data are the mean ± SEM of three independent experiments. (E) Lysates from MDCK-SIAT cells infected with the indicated viruses at high MOI and harvested at 16 h p.i. were analysed by SDS-PAGE and western blotting for the indicated polypeptides.

Given the high proportion (97.9%) of pdm2009 isolates encoding uAUG (Table S1), we analysed three clinical isolates with known, limited *in vitro* passage histories (29) by western blotting following high multiplicity infection of MDCK-SIAT cells. All three viruses produced both NP and eNP (Fig 4E), further supporting the potential *in vivo* relevance of eNP expression.

The terminal regions of IAV genome segments are involved in vRNA packaging, via specific RNA signals (30). To test whether the mutations that added or subtracted the uAUG codon affected segment 5 packaging, RNA was extracted from independently grown stocks of WT or mutant viruses and the amount of segment 5 measured by qRT-PCR. The values obtained were then considered as a ratio to the plaque titre of the stocks, normalised to WT PR8, to produce a relative vRNA:PFU ratio. As a positive control for a virus with a packaging defect, we analysed a PR8 mutant with two clusters of synonymous mutations (9 nucleotide changes in total) introduced into the 5’-ends of segments 4 and 6, where bioinformatics analyses had predicted the likely location of packaging signals (31). The genome copy:PFU ratio of this “4c6c” virus was elevated by over 2 log_10_ compared to WT virus (Fig 5A). In contrast, introduction of the C28A mutation into PR8 had very little effect on the quantity of segment 5 required to form an infectious unit. Replacement of PR8 segment 5 with the corresponding WT Eng195 vRNA elevated the genome:PFU ratio by ∼ 10-fold, suggestive of a packaging incompatibility between the PR8 backbone and the pdm2009 segment. However, addition of the A28C mutation into the Eng195 segment to remove the uAUG codon did not worsen this phenotype. Overall therefore, the data did not indicate any large effect of the A28C polymorphism on segment packaging.

**Figure 5.**
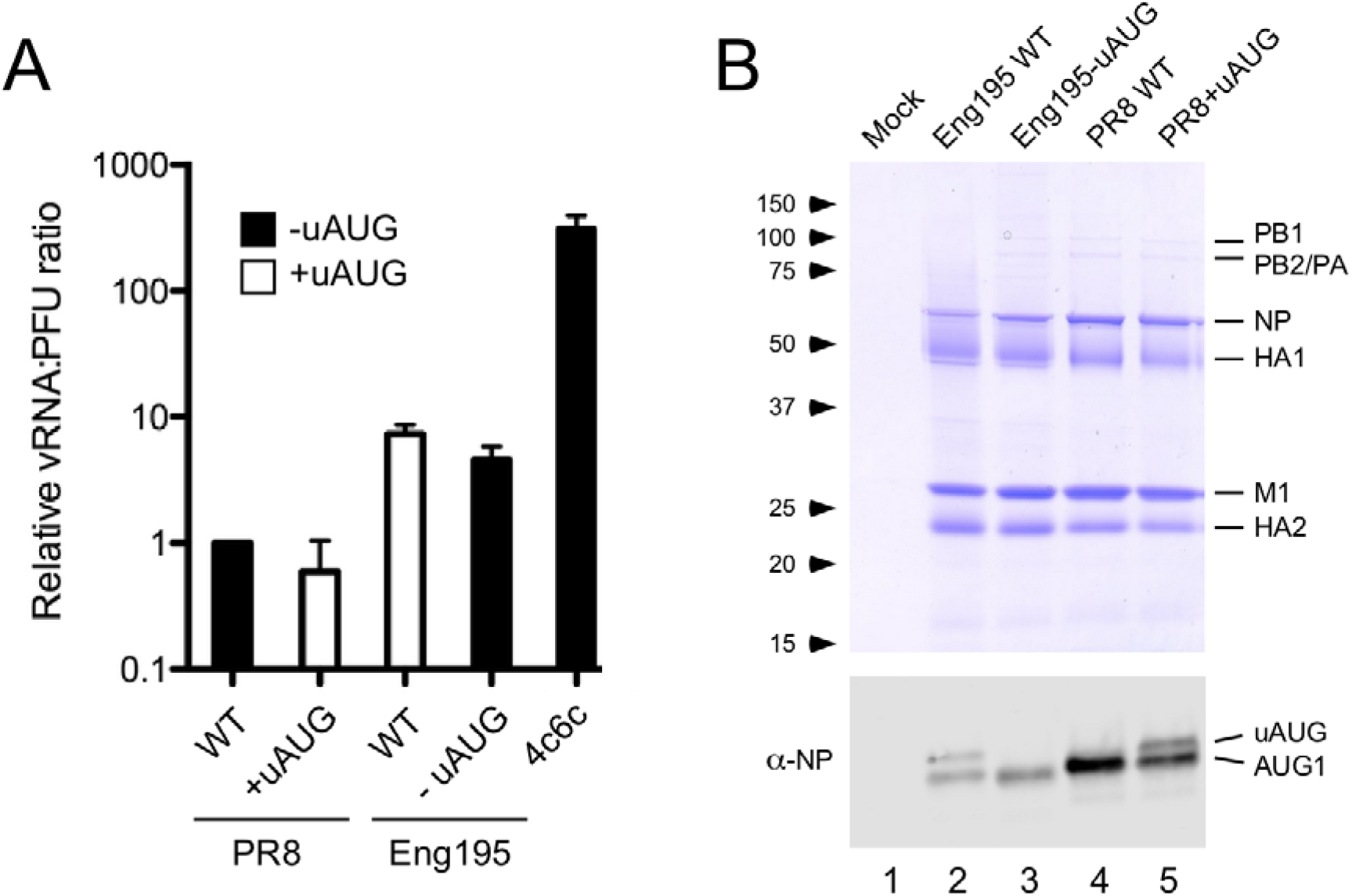
Virion composition of PR8 and PR8 7:1 reassortants containing the Eng195 segment 5 (Eng195). (A) The segment 5 vRNA content of virus stocks of known infectious titre was determined by qRT-PCR and the values used to derive genome:PFU ratios, normalised to that of WT PR8. Data are the mean ± range of two independent replicates. (B) Aliquots of sucrose gradient purified virus or the corresponding fraction from uninfected allantoic fluid was analysed by SDS-PAGE and (upper panel) Coomassie Blue staining or (lower panel) western blotting for NP. The migration positions of molecular mass standards (kDa) and major viral structural proteins are indicated. Note that the gel used for the lower panel was run further to separate the two NP isoforms.

To determine if eNP is incorporated into virus particles, virus stocks were grown in embryonated hens’ eggs and virions purified from allantoic fluid by pelleting through a sucrose cushion followed by banding on a sucrose velocity gradient. SDS-PAGE and Coomassie blue staining of the resulting material showed the presence of the expected major viral structural proteins (Fig 5B, top panel). Re-analysis of the same material by western blotting for NP under PAGE conditions sufficient to separate the two forms of NP clearly showed the presence of both canonical and eNP in an approximately 2:1 ratio in viruses where the uAUG codon was present (Fig 5B, bottom panel). Thus consistent with its apparently normal function in minireplicon assays, eNP was incorporated into virus particles.

### The C28A polymorphism influences pathogenesis in mice

To examine the role of eNP *in vivo*, the mouse model of IAV infection was used. First, groups of BALB/c mice were infected with PR8 or the PR8:Eng195 segment 5 reassortant in either WT form or with the C28A polymorphism, and weight loss followed over 5 days. Uninfected mice gained weight over time, while mice infected with WT PR8 lost around 5% of their starting body weight (Fig 6A). Unexpectedly, the PR8 C28A mutant induced significantly greater weight loss in the animals, resulting in an average loss of over 15%. Consistent with this, the PR8 C28A-infected mice showed increased clinical signs compared to their WT-infected counterparts, including increased respiratory rate, lower motility, more extreme staring of the coat and more emphatic hunching (data not shown). Animals infected with the PR8:Eng195 segment 5 WT or A28C viruses did not show obvious clinical signs or lose any substantial amount of weight over the 5 days (Fig 6A). At day 5, all animals were sacrificed and the lungs collected for further analyses. When viral loads were measured, both WT and C28A PR8 viruses gave titres of around 10^6^ PFU/ml of homogenate (Fig 6B). The reassortant virus with WT Eng195 segment 5 produced titres of around 10^5^ PFU/ml, despite the lack of clinical signs of infection. However, the corresponding A28C mutant lacking the uAUG codon gave substantially lower (on average, almost 2 log_10_) titres, suggesting attenuated virus replication. Examination of lung homogenates by western blotting for viral NP confirmed that the PR8 C28A virus expressed eNP *in vivo* (Fig 6C). Neither form of NP could be detected in material from animals infected with the PR8:Eng195 reassortant viruses, most likely because of the lower levels of virus replication. To measure innate immune response stimulation, levels of IFN-ß mRNA in the lung homogenates were assessed by qRT-PCR. Transcripts were undetectable from mock infected mouse lung but were clearly induced by PR8 virus infection (Fig 6D). However, despite the more severe disease seen with the PR8 C28A virus, there was no significant difference between this and WT virus samples. IFN-ß mRNA levels were substantially lower in animals infected with the WT PR8:Eng195 virus and were undetectable in all but one animal infected with the A28C mutant (Fig 6D); these differences plausibly correlated with virus load (Fig 6B). A similar outcome was obtained when a broader array of cytokines and chemokines were analysed; few differences of note between the PR8 pair of viruses and generally higher induction from WT Eng195 than its A28C counterpart (Fig S6).

**Figure 6.**
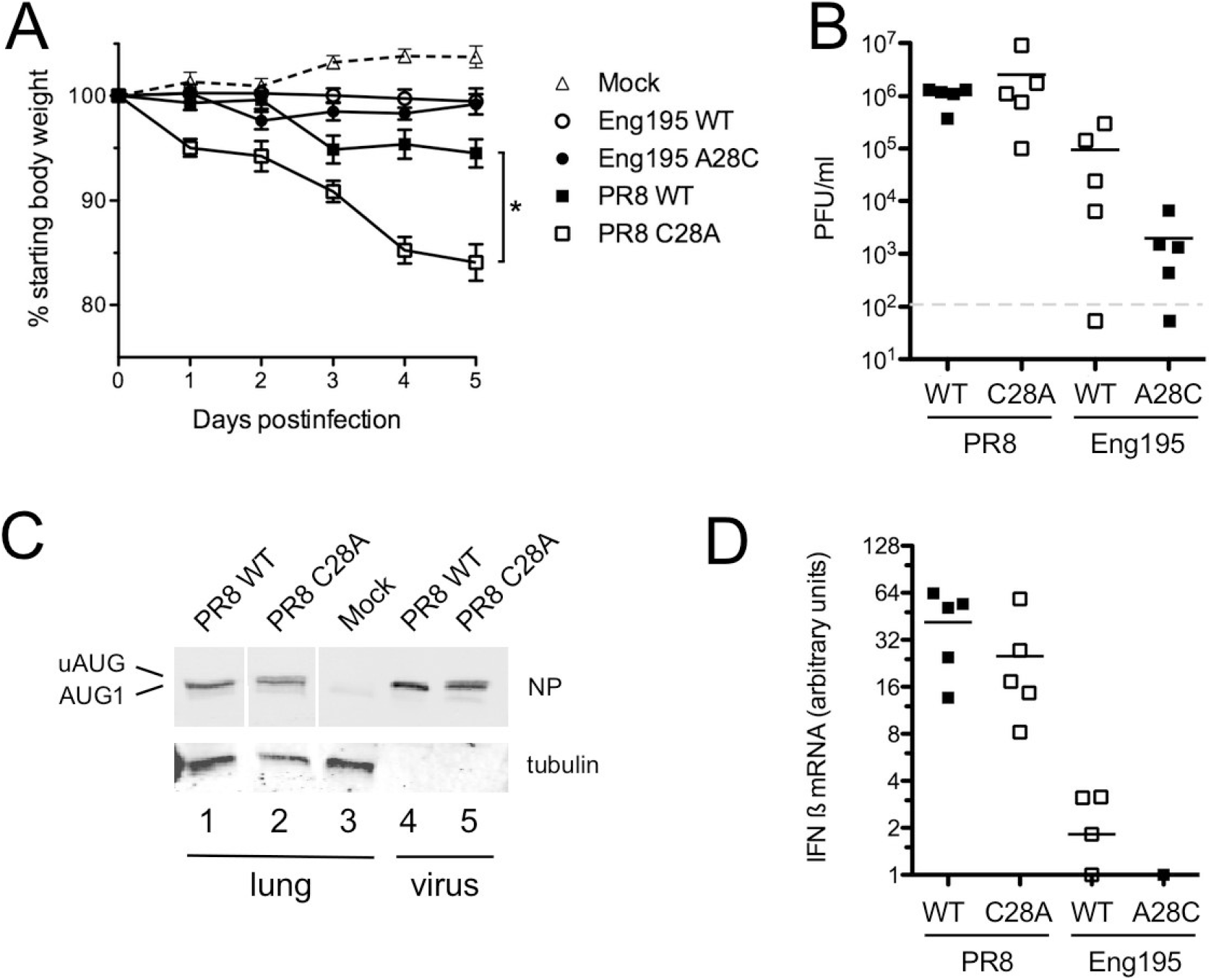
Pathogenesis of eNP-expressing viruses in Balb/c mice. Groups of 5 six-week old mice were infected with 200 PFU or PR8 viruses or 500 PFU of 7:1 PR8 reassortant viruses containing the Eng195 segment 5 (Eng195). (A) Body weight was measured daily for 5 days after infection. Data are plotted as the mean ± SEM. * *p* < 0.05 (One way ANOVA with Tukey’s multiple comparison post test; WT PR8 versus PR8 C28A). (B) Animals were euthanised at day 5, the left lungs homogenised and virus titres determined. Dashed line indicates limit of detection. Differences between pairs of viruses were non-significant, as assessed by *t-*tests. (C) Aliquots of pooled lung homogenate (lung) or purified virus (virus) were analysed by SDS-PAGE and western blotting for the indicated proteins. (D) RNA was extracted from the left lung tip and IFN ß and GAPDH mRNA levels determined by qRT-PCR. IFN ß transcript was not detected in RNA from uninfected animals, so positive values were corrected for GAPDH levels and then expressed relative to the lowest samples that gave a C_t_ value (one animal each from WT and A28C Eng195 infections). Differences between virus pairs were not statistically significant (non-parametric *t*-tests).

To assess histopathological changes in the mice, formalin-fixed lung sections were stained with haematoxylin and eosin and examined by a veterinary pathologist. Changes identified in infected mice were consistent with acute to subacute IAV infection; these were characterised by degeneration and necrosis of epithelial cells lining airways, accompanied by peribronchial and perivascular inflammation, as well as interstitial inflammation and necrosis (Fig 7A and data not shown). The inflammatory infiltrate consisted of lymphocytes and macrophages with fewer plasma cells, and rare neutrophils and eosinophils. When the slides were scored blind for various pathological features, the C28A PR8 mutant gave generally higher scores than WT PR8 in most categories (Fig 7B). Combining these scores along with a consideration of the area of lung affected by pathological changes to give an overall score showed significantly higher (*p* < 0.05, *t*-test) damage from the PR8 C28A virus (Table S5). Conversely, the A28C Eng195 mutant gave lower average scores in all categories than its WT counterpart and an overall highly significant difference of *p* < 0.005 (Fig 7C and Table S4), confirming that mutation of the uAUG codon substantially attenuated virus pathogenicity.

**Figure 7.**
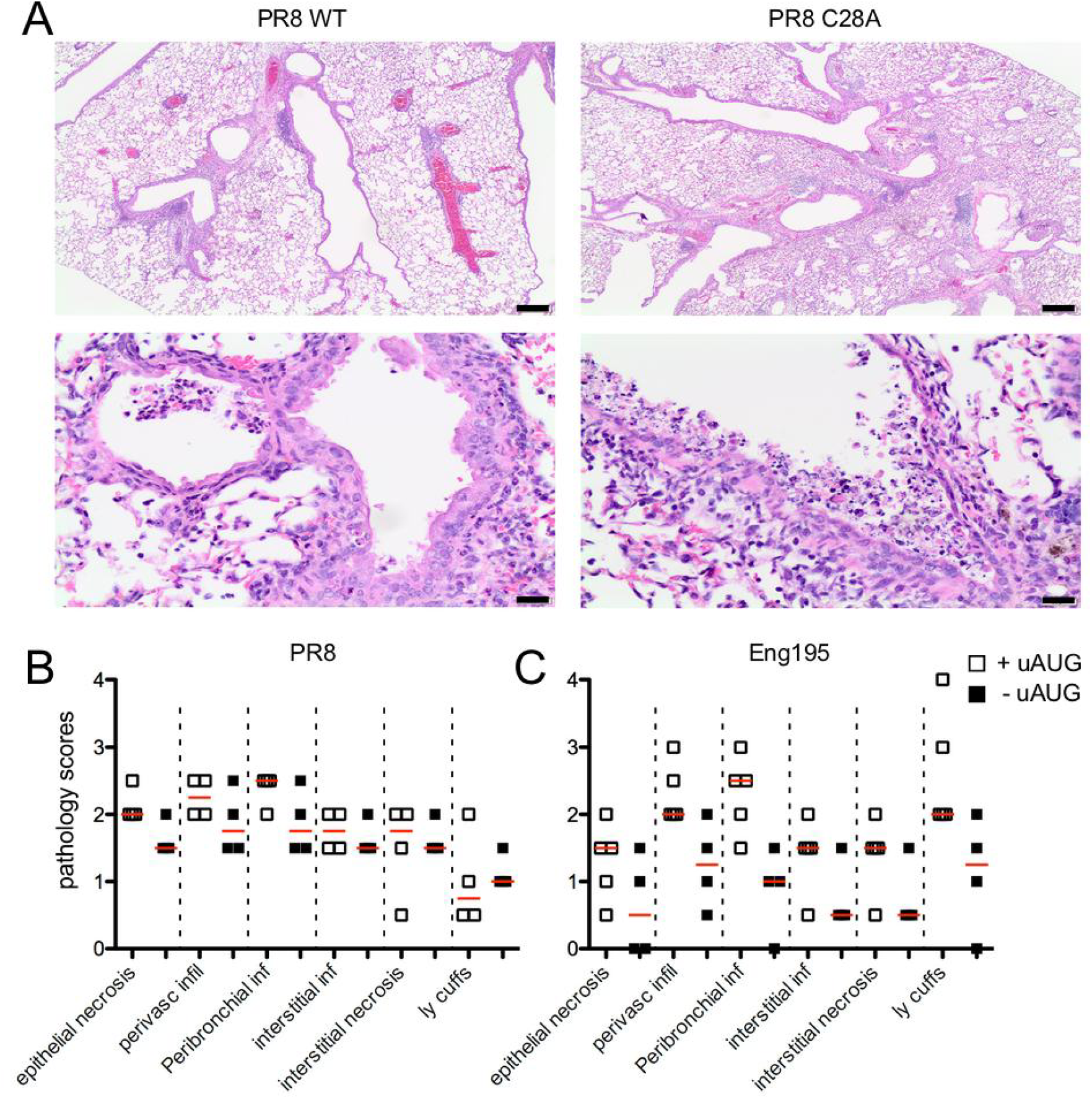
Histopathology of eNP-expressing viruses in Balb/c mice. At day 5 p.i., the right lung lobes of inoculated mice were collected, fixed, processed, and (A) stained with hematoxylin and eosin. Mock-infected mice showed no significant pathology (Table S4). Scale bars indicate 200 µm (top panels) or 20 µm (lower panels). (B, C) The severity of the pathology in individual lungs was assessed in a blind manner, and an overall score out of 4 for the various categories of damage was assigned (infil; infiltrate, inf; inflammation, ly; lymphocyte). Red bars indicate the median.

BALB/c mice are biased towards Th2-type responses (32) and mouse strain-dependent variations in response to pdm2009 infection have been observed (33). Accordingly, to further test the effect of modulating NP start codons on viral pathogenicity, we examined the course of infection in the outbred CD-1 mouse strain after infection with a further two pairs of recombinant viruses differing only in the presence or absence of the segment 5 uAUG codon: a complete clone of the SW210 pdm2009 virus, and the St Jude Children’s Hospital clone of PR8 (34). Infection with both WT and C28A mutant PR8 viruses led to severe weight loss from day 3 post infection onwards, resulting in all animals reaching a humane endpoint by day 11. However, consistent with the previous experiment, animals infected with the eNP-expressing mutant version of PR8 lost weight faster and died sooner (Fig 8A). Infection with WT SW210 virus led to animals losing around 10% of their body weight by day 7 followed by recovery from day 10. In contrast, the A28C derivative did not cause any evident disease (Fig 8A). Examination of lung titres taken at days 3 and 5 p.i. confirmed that the animals were infected but that the WT SW210 virus had replicated to titres over 1 log_10_ higher than the A28C mutant (Fig 8B), indicating that removal of the NP uAUG codon was attenuating *in vivo* in the background of an authentic pdm2009 virus. Thus, the attenuating effect of altering the uAUG codon was consistent across virus strains and breeds of mice.

**Figure 8.**
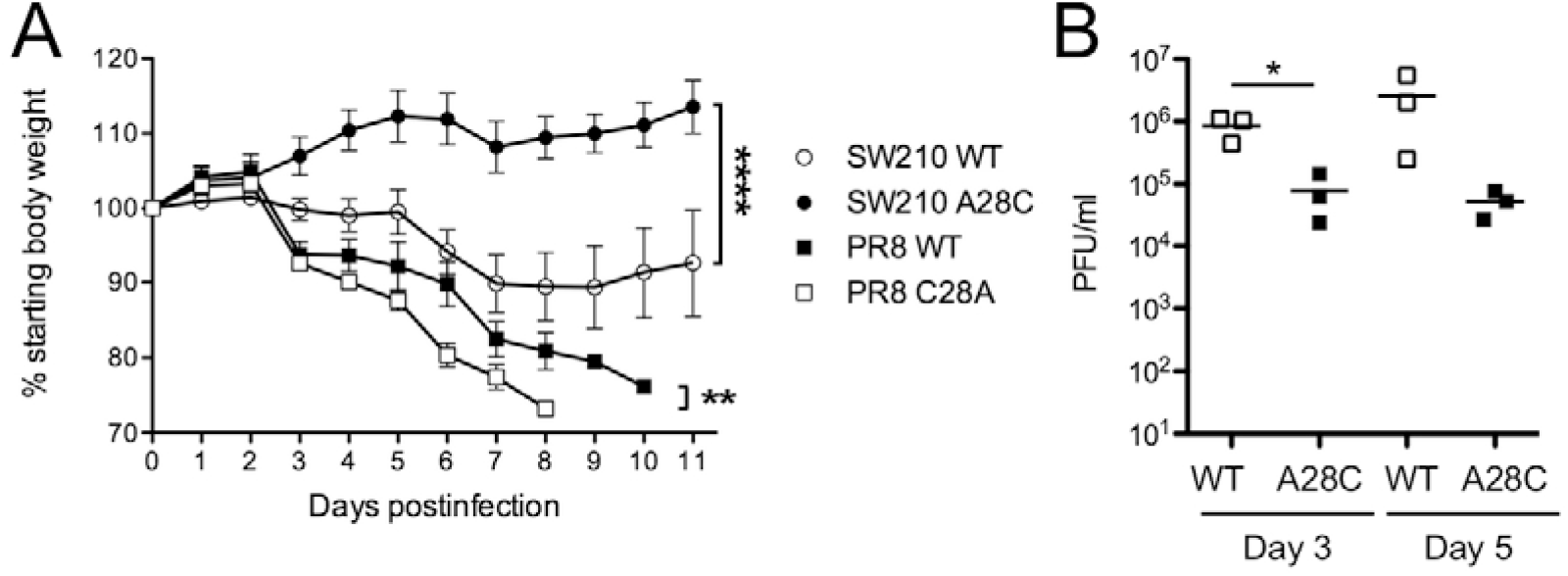
Pathogenesis of eNP-expressing viruses in CD-1 mice. Groups of 4-5 animals were infected with 10^4^ PFU of PR8 viruses or 5×10^5^ PFU of SW210 viruses. (A) Body weight was measured daily for up to 11 days. Animals that met the humane end-point were euthanized earlier. Data are the mean ± SEM. ** *p* < 0.01, **** *p* < 0.0001 (*t*-tests between pairs of viruses). (B) For the SW210 pair of viruses, an additional six animals were included, and three animals were euthanized on each of days 3 and 5 p.i. for virus titres in lung homogenates to be determined. *p* < 0.05 (*t-*test between pairs of viruses on separate days).

### The C28A polymorphism influences pathogenesis in pigs

With evidence from the mouse model of IAV infection that the presence of the segment 5 uAUG increased virulence, we tested whether this phenotype was replicated in pigs, where uAUG appeared to be strongly selected for in an evolutionary context. For this, we utilised a previously characterised challenge system using Babraham inbred pigs (35,36) for the Eng195 virus. Groups of animals were infected intranasally with 2.2 x 10^5^ PFU of WT or A28C Eng195 and monitored for virus shedding by daily nasal swabs for 4 days. All animals shed detectable levels of virus for the duration of the experiment and although the average titres were higher from animals infected with the A28C virus at days 3 and 4 p.i. (Fig 9A), the data were variable and the differences were not statistically significant. At day 4 p.i., animals were euthanized and samples taken from the respiratory tract for virus titration. Titres were highest in tracheal swabs, intermediate in bronchiolar lavage fluid (BALF) and lowest in lung tissue homogenates, where not all samples were detectably positive (Fig 9B). As with the shedding data, there were no significant differences between the two viruses however. Examination of the animals’ lungs showed areas of interstitial pneumonia and atelectasis mostly in the apical lung lobes (Fig S7). However, the overall macroscopic pathology scores between the two groups were also not significantly different (Fig 9C). To examine microscopic pathology, five tissue samples per right lung (two apical and one each from the medial, diaphragmatic and accessory lobes) were formalin fixed, processed into paraffin-wax and cut sections stained with H&E. Histopathological analysis showed multifocal interstitial pneumonia, attenuation/necrosis of bronchial and bronchiolar epithelial cells, presence of inflammatory cell infiltrates within the interalveolar septa and the alveolar lumen, and oedema (Fig 10A, B). These histopathological changes were scored across all sections by a board-certified veterinary pathologist according to five parameters: necrosis of the bronchiolar epithelium, airway inflammation, perivascular/bronchiolar cuffing, alveolar exudates, and septal inflammation (Table S6). Here, a clear difference between the two viruses became apparent, with the A28C virus on average provoking lesser amounts of damage to the lung by each criterion (Fig 9D). To assess virus spread within the lung, sections were stained by IHC for IAV NP. Viral NP was observed mainly within the bronchial and bronchiolar epithelial cells (Figure 10C), but also within inflammatory cells inflitrating into the bronchiolar lumen and alveolar spaces. NP-IHC staining was scored for the numbers of antigen-positive cells in airway epithelia and alveolar septa/lumens, showing reduced numbers of cells infected with the A28C mutant (Fig 9E). When histopathological and IHC scores were combined to provide an overall measure of pathology (the “Iowa” scale) (35), animals infected with the WT virus had consistent and significantly worse disease than those infected with the A28C mutant virus (Fig 9F). Thus, removing the uAUG codon from a pdm2009 virus led to reduced virulence in a biologically relevant large animal model of IAV infection.

**Figure 9.**
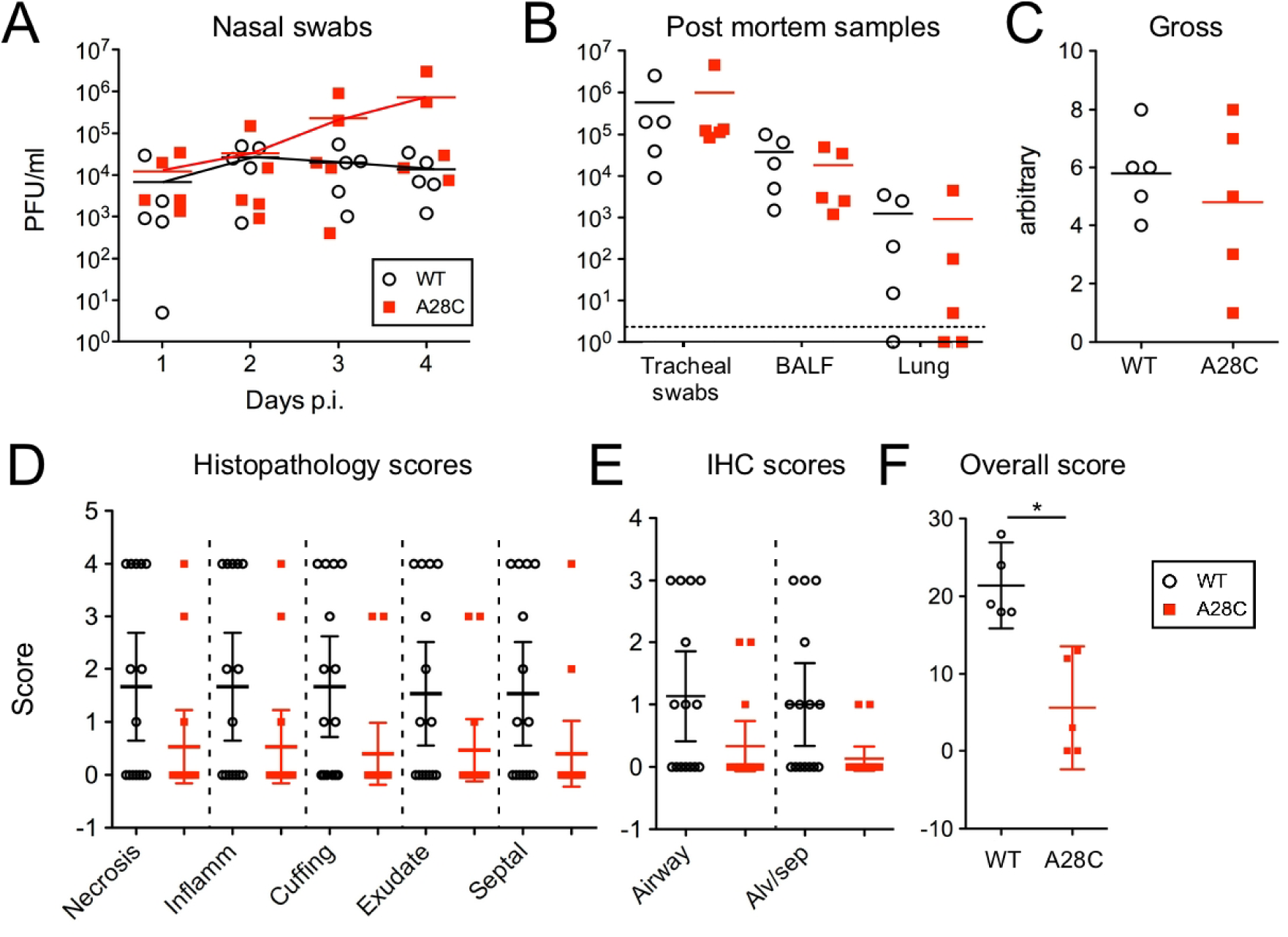
Pathogenesis of a pdm2009 virus with altered eNP expression in pigs. Inbred Babraham pigs were challenged with WT or A28C Eng194 and (A, B) swabs and samples taken as indicated and titred for virus. Dashed line indicates the limit of detection. (C) Following necropsy at day 4 p.i., lungs were removed and scored for gross pathology. (D, E) Cut tissue sections were blinded and (D), stained with H&E and scored for the indicated categories of pathology or (E), stained for NP and scored for the quantity of viral antigen-positive cells. (F) – Iowa overall score, taking D and E together. * = *p <* 0.05 (Mann Whitney *t*-test).

**Figure 10.**
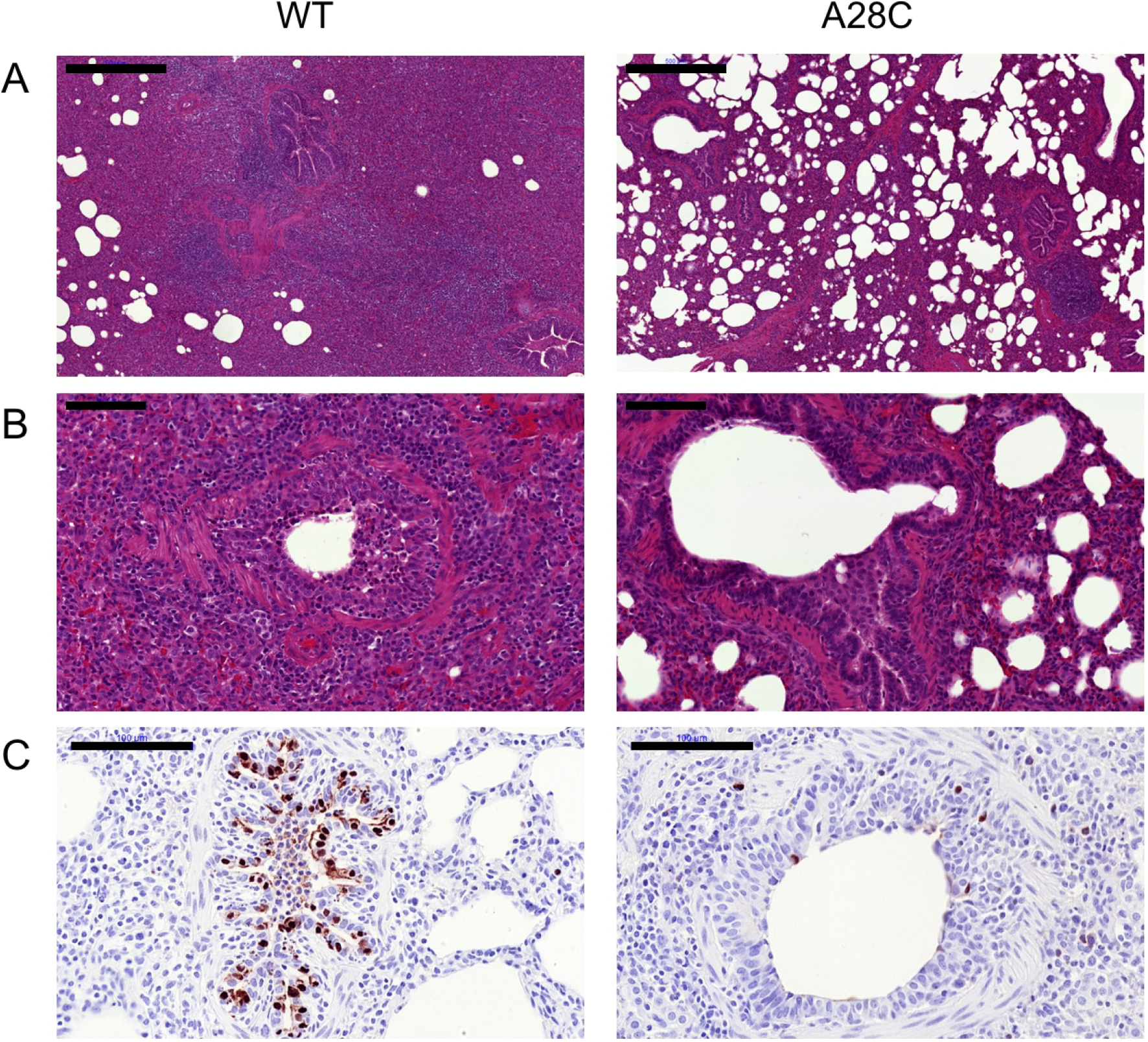
Histopathological analyses of pig lungs 4 days post challenge with WT and A28C Eng195 viruses. Representative sections of lung stained with (A, B) H&E and imaged at low (scale bar = 500 µm) and high (scale bar = 100 µm) magnification respectively or (C) stained for IAV NP (in brown) and counterstained with haematoxylin. Scale bar = 100 µm.

## Discussion

The pdm2009 virus possesses several genetic features which might have explained its unexpectedly mild disease characteristics in humans, including an avian IAV-like signature at PB2 residue 627 and truncated PB1-F2, PA-X and NS1 genes. However, artificially altering these sequences to what could reasonably be predicted to be a more pathogenic form and testing them in animal models of infection has generally failed to support a causative role in disease attenuation (6,37-43). Here, we investigated another genetic quirk of the pdm2009 virus; the presence of an extra in-frame start codon in the 5’-UTR of the NP gene. We found that this uAUG originally emerged in classical swine viruses circulating in the early 1960s, before being inherited by various reassortant lineages of viruses including the pdm2009 strain. We showed that the uAUG codon is used for translation initiation to produce two isoforms of NP in infected cells, in a roughly 3:1 ratio of “normal” and “extended” [eNP] polypeptides. This had little apparent functional consequence *in vitro*, either for viral gene expression (something that NP plays a crucial role in) or for overall virus replication. However, while the presence or absence of the uAUG did affect pathogenicity in mice and in pigs, it acted to increase rather than decrease virulence in both animal models of infection. Importantly, this included pigs, where the human pandemic strain originated. Thus, once again, investigation of an unusual genetic feature of the pdm2009 virus has not supported the simple hypothesis of an attenuating role in virulence.

A previous study has examined the functional significance of the segment 5 uAUG; the authors did not directly determine whether it was actually used for translation initiation, but using RNP reconstitution assays, they concluded that its presence significantly influenced viral transcriptional activity, albeit only by around 2-fold (7). Here, we saw similar magnitude effects, but without achieving statistical significance (Fig 2). We consider such small fluctuations in minireplicon activity unlikely to have much biological consequence; viruses with the same mutations replicated similarly in a variety of *in vitro* settings and had no obvious deficits in gene expression (Fig 3). The question therefore remains, of how the presence of the uAUG might enhance virulence. Whilst it is difficult to rule out subtle effects arising directly from the UTR mutation (*e.g.* on segment 5 RNA synthesis or packaging), these were normal as far as we could measure (Figs 4, 5). Instead, we prefer the hypothesis that the *in vivo* phenotypic change results from the expression of a new isoform of NP. The question then arises of how the function of the novel polypeptide varies from that of NP.

The N-terminal 20 amino acids of the canonical form of NP appear to be flexible, as they are not visible in crystal structures of the whole protein (11,12,44). The primary function attributed to this region of NP is that of a non-classical nuclear localization signal (ncNLS) that binds cellular importin α (14,45-47). It is therefore possible that the addition of the 6 amino acids unique to the eNP sequence could affect interactions with importin α; the much larger (50 amino acid) extension on the related influenza B virus NP has been shown to affect nuclear localization of the protein (48). However, the structure of the IAV NP ncNLS bound to importin α suggests that this would not necessarily be the case here, as it binds to the relatively shallow minor NLS-binding site, leaving the N-terminus of the NP sequence free, where a short extension could easily be accommodated (47). In addition, we did not see any appreciable differences in NP intracellular trafficking arising from the presence or absence of eNP during a time course of infection in A549 cells (data not shown). The N-terminal disordered region of NP is also a target for regulatory post-translational modification, such as phosphorylation (49-51) and sumoylation (52); again something that could potentially be affected by the addition of the 6 amino acids unique to eNP. However, the post translational modifications only occur on a small fraction of the NP molecules in the cell and would presumably still occur normally on canonical NP, which remains the most abundant isoform in cells infected with a virus containing the uAUG codon. Thus, any changes to post translational modification of eNP seem unlikely to be the primary cause of the phenotypic effects seen here.

Our working hypothesis is therefore that acquisition of the segment 5 uAUG codon represents a gain-of-function mutation for porcine IAV, reflecting a novel function for eNP versus NP. Since NP has not been convincingly associated with any intrinsic enzymatic activity, this would most simply be explained by the additional sequence mediating a new (or stronger) interaction with a cellular binding partner that affects the outcome of *in vivo* infection, but without affecting virus replication *in vitro*, at least in the cell lines tested. At present, we do not have candidates for such a cellular factor and further experimentation is required to identify these. Another question that remains to be answered is whether eNP modulates pathogenesis through incorporation into viral RNPs, or as an isolated protein.

We found through evolutionary analyses that the segment 5 uAUG is primarily a trait associated with virus strains of swine origin, where its recurrent emergence and fixation in the major swine virus lineage implies it provides a host-specific selective advantage. This hypothesis is consistent with the reduced virulence exhibited by pdm2009 virus engineered to lack the uAUG codon (Figs 9, 10). Whether the uAUG codon provides a selective advantage in the human host is also unclear, but the sporadic emergence of the uAUG in humans prior to 2009 without provoking a selective sweep implies these events, and its dissemination in 2009, were driven by founder effect. However, its subsequent maintenance in the human pdm2009 lineage (Table S2) gives no sign that it is being selected against in humans.

In summary, we have characterised a swine host-specific mutation in the 5’UTR of IAV segment 5 which introduces an alternative start codon in frame with the NP ORF that adds 6 amino acids to the N-terminus of the protein. This mutation modulates virulence in mice and pigs, and was introduced to the human population via the 2009 pandemic. This knowledge adds to our ability to understand and predict IAV virulence in specific hosts and furthermore, suggests that the tendency for IAV sequencing efforts to disregard the viral UTRs may be missing useful information.

## Materials and Methods

### Bioinformatics

On 31 Jan 2019, all full-length NP nucleotide sequences of IAV (any host, any location, any year) were downloaded from Genbank via the NCBI Influenza Virus Resource database (53). The sequences were named using the schema: “>{serotype}_{host}_{accession}_{strain}_{country}_{year}/{month}/{day}_{segname}”. 50471 sequences in total were downloaded and of these there were 48288 with known subtype, host, country and year. These were screened for duplicates and quality (number of ambiguities), leaving 48245 good quality sequences which were padded with 10 codons (30 nucleotides) and then roughly aligned for further processing. However, although these sequences were tagged as ‘full length’, not all of them reported sequence before the usual NP start codon, leaving 33622 which had 5’UTR data suitable for analysis, representing 69.7% of the good quality sequences. In order to determine in which major lineages the sequences with upstream start codons occur, a stratified subsampling was performed, based on a maximum of 1 sample per joint category of host + subtype + country or state (if USA, Canada, China or Russia) + year + upstream start codon. This type of subsampling retains the diversity of hosts, subtypes, locations, dates and start codons but will skew the percentages of hosts vs start codons. The subsampling resulted in a data set of 6242 sequences, which were aligned using MUSCLE in MEGA and manually adjusted (at the far 3’ end of the coding region which was not guaranteed to be complete by the process).

To construct phylogenetic trees, phylogeny was calculated from the stratified subsample of 6242 sequences using RAxML with the GTR model allowing for a gamma distribution of variable site rates and 100 bootstraps. Detailed time-resolved phylogenetic trees of selected lineages or clades of the 6242 sequences dataset were inferred with BEAST (1.10.4) (54) using the SRD06 codon partitioned nucleotide model with uncorrelated relaxed log normal clock models and the constant population size or skygrid tree priors. Per lineage or clade, 1000 trees were sampled from the resulting posterior distribution of trees (after 10% burn-in). For each selected clade or lineage, the start codon was mapped as a binary variable (Conventional or Upstream) onto the set of 1000 posterior trees using a discrete trait asymmetric model in BEAST, resulting in a set of 10,000 trees (approximately 10 mappings per original tree), which were summarised to a maximum clade credibility tree using TreeAnnotator

### Plasmids and antisera

Each of the eight IAV segments used throughout were encoded on IAV reverse gene plasmids containing bi-directional RNA polymerase I and II promoters. Two different PR8 strains were used: most experiments used an MDCK-adapted variant of the UK National Institute of Biological Standards and Control vaccine strain of PR8 (55), while data shown in figure 7 used an egg-adapted PR8 strain described previously (28,34). Plasmids for the A/England/195/2009 (Eng/195; an early UK pdm09 strain) are described in (22), while those for A/Halifax/210/2009 (SW210; an early pandemic isolate from the Queen Elizabeth II Hospital, Halifax, Nova Scotia, Canada) are described in (56). A plasmid encoding a gene fusion between the 5‘-201 nucleotides of PR8 segment 5 and GFP was made by PCR-cloning the appropriate IAV sequence into pEGFP-1 (Clontech), followed by oligonucleotide-directed PCR mutagenesis (using standard protocols) to remove the A of the GFP AUG codon. Nucleotide substitutions in the segment 5 plasmids to modify AUG motifs were made by further rounds of oligonucleotide-directed mutagenesis. To create the 4c6c PR8 virus with a genome packaging defect, codons 546-548 of the PR8 HA gene were synonymously mutated to **t**T**a**GG**t**GC**c** and codons 451 to 453 of PR8 NA were mutated to **tca**AT**a**GA**t** (altered nucleotides indicated in lower case bold, all sequences given in (+) sense). Primer sequences are available on request. Sequence modifications to plasmids were confirmed by commercial Sanger sequencing (GATC, Eurofins Genomics) before being used in downstream experiments.

For western blotting, purchased antisera used were: mouse monoclonal anti-GFP (clone JL8, Clontech), rat monoclonal anti-ß tubulin (clone YL1/2, Abd-Serotech) and mouse monoclonal anti-IAV NP (clone AA5H, Abd-Serotech). In-house rabbit polyclonal antisera raised against IAV PB1, PB2, NP and M1 have been previously described (57-59). Secondary antibodies labelled with infrared-fluorescent dyes were obtained from Fisher. For IHC staining of cut tissue sections, mouse monoclonal anti-NP (hybridoma HB-65 from ATCC (Manassas, VA, USA)) was used.

### Viruses

Virus rescues were performed as previously described (27,60). Briefly, 293T cells were transfected with 250ng each of the 8 plasmids from a viral reverse genetics set (or no segment 5 as a negative control), with 1µg/ml tosyl phenylalanyl chloromethyl ketone (TPCK)-treated trypsin added at 48 hours post-transfection. At 72-96 hours post-transfection, supernatants were harvested and either passaged on MDCK cells in serum free medium with 1µg/ml TPCK-treated trypsin, or 100 µl was inoculated into the allantoic cavity of 12 day-old embryonated hen’s eggs, the allantoic fluid of which was harvested at day 14 and aliquoted as virus stock. Titres were determined by plaque assay on MDCK (PR8; at 37°C) or MDCK-SIAT cells (61)(Eng/195; at 35°C). Plaques on MDCK cells were typically visualized by toluidine blue staining, while plaques on MDCK-SIAT cells were immunostained for viral NP. The presence of the desired mutations in the virus genome was confirmed by RT-PCR and Sanger sequencing of RNA isolated from the virus stocks. Early-passage H1N1 pdm2009 clinical isolates A/Nottingham/Adult-Community04/2009 (AC04), A/Nottingham/Child-Community06/2009 (CC06) and A/Nottingham/Child-Community07/2009 (CC07) were isolated and passaged twice on MDCK cells as described (29).

### Cells and transfection methods

293T, MDCK, A549, and NPTr cells were grown in DMEM supplemented with 10% FBS and 1% penicillin and streptomycin (Fisher) and maintained by twice weekly passage. IAV minireplicon assays were performed as described (27). Briefly, unless otherwise stated, 50ng each of pDUAL plasmids encoding segment 1,2, 3 and 5 and 20ng of a construct that expresses an IAV-like vRNA encoding luciferase were transfected into 293T cells in 24 well format. Transfected cells were harvested in 100 µl cell culture lysis reagent (Promega) and 60 µl supernatants were mixed with 25 µl 6mM beetle luciferin (Promega). Luminescence was measured on a GloMax luminometer (Promega). Assays were performed using four technical replicates for each datapoint as well as at least three biological repeats.

### Protein methods

*In vitro* translation reactions were performed using coupled bacteriophage T7 RNA polymerase transcription-rabbit reticulocyte lysate translation reactions as per the manufacturer’s instructions (Promega TNT). Samples were radiolabelled with ^35^S-methionine (Perkin Elmer) and detected by SDS-PAGE and autoradiography. To separate eNP and NP, samples were loaded onto 10% pre-cast gels (BioRad) and run until the 50kDa ladder marker had just run off the bottom of the gel. For western blotting, wet transfers were performed, membranes were blocked with 5% milk for 30-60 minutes then incubated with specific antibodies in 2% BSA at 4°C overnight. The next day, membranes were incubated with secondary anti-rabbit or anti-mouse IgG antibodies conjugated to fluorophore AlexaFluor 680 or 800 as required, before imaging using a LiCor Odyssey FC.

To purify virus, allantoic fluid was clarified twice by centrifugation for 10 min at 2100 x g, then loaded onto a 30% sucrose/PBS cushion and spun at 30,000 rpm using an SW28Ti Beckman rotor for 1 hour and 30 min at 4°C. The resulting pellet was gently washed once with 500 μl of PBS to remove residual sucrose and re-suspended back in 50 μl of PBS overnight. The virus was further purified by ultracentrifugation through a 15-60% sucrose gradient (in PBS) spun at 38,000rpm for 40 min using a Beckman SA41Ti rotor at 4°C. The virus band was extracted from the gradient using a syringe and virus pelleted by centrifugation at 30,000 rpm for 90 min at 4°C before being resuspended as above.

For cytokine arrays of mouse lung homogenates, 20 µl of lung homogenate per mouse was collected and pooled within groups. Cytokines were then measured using Proteome Profiler Mouse Cytokine Array Kit (R&D Biosystems) according to manufacturer’s instructions. Arrays were imaged and spot intensities quantified using a LI-COR Odyssey Infrared Imaging System (LI-COR, Cambridge, UK). Following normalization to within-assay control spots values were plotted as fold increase in cytokine expression over mock-infected animals.

### Ethics statement

Animal experimentation was approved by the Roslin Institute Animal Welfare and Ethical Review Board, the Pirbright Institute Ethical Review Board under the authority of Home Office project licences (60/4479 and 70/7505 respectively) within the terms and conditions of the UK Home Office “Animals (Scientific Procedures) Act 1986“ and associated guidelines, or in compliance with the guidelines of the Canadian Council on Animal Care (CCAC) as outlined in the Care and Use of Experimental Animals, Vol. 1, 2nd Edn. (1993). In this case, the animal care protocol was approved by the University of Ottawa Animal Care Committee (Protocol Number: BMI-85) and all efforts were made to minimize suffering and mice were euthanized at humane end-points, if infection resulted in greater than 30% body weight loss plus respiratory distress.

### Mouse infections

BALB/c mice were purchased from Harlan UK Ltd (Oxon, UK), CD-1 mice were purchased from Charles River Laboratories, (Montreal, Quebec, Canada). Five- to 12-week-old female mice were used in all experiments. A group size of 5 was used as based on variance observed in previous experiments, this was expected to give 80% power to detect a statistically significant difference of 6% weight loss at the 5% significance level. Mice were anaesthetized using isoflurane (Merial Animal Health Ltd) and infected intranasally with virus in 40 μl serum-free DMEM. Mice were weighed daily and assessed for visual signs of clinical disease, including inactivity, ruffled fur and laboured breathing. At day 5-post infection, mice were euthanized by CO_2_ asphyxiation. Virus titration and RNA extraction and qRT-PCR was undertaken as previously described (62). Briefly, left lung homogenates were collected in 500 µl DMEM and homogenised using a Qiagen TissueLyser II. For virus titration, standard plaque assays on MDCK cells were performed and the remaining supernatant was used for RNA quantification. RNA was extracted using a Qiagen Viral RNA mini kit according to manufacturer’s instructions and DNAse treated using Promega RQ1-RNAse free DNAse. RT-qPCR was undertaken using a BioLine Sensifast one step RT-qPCR kit with modified cycling conditions of 45°C for 10 minutes, 95°C for 2 minutes, then 40 cycles of 95°C for 10s and 60°C for 30s. Primer sequences are given in Table 2. For histopathological analysis, the four lobes of the right lung were inflated with and then immersed in 10% neutral buffered formalin (Sigma-Aldrich) until fixed, then processed using routine methods and embedded in paraffin blocks. 5 µm thin section were then cut and stained with haematoxylin and eosin for histological examination. Sections were assessed (blinded) by a veterinary pathologist (PMB). The individual pathology features assessed were damage to the airway epithelium (degeneration, necrosis and repair), perivascular inflammation, peribronchi/bronchiolar inflammation, interstitial inflammation, interstitial necrosis and type II pneumocyte hyperplasia. Each feature was scored from 1 (mild) to 3 (marked). The percentage of lung affected was also noted.

### Pig infection model

Ten 12-14 wk old Babraham large white inbred pigs (average weight 30 kg) were obtained from the Pirbright Institute/ Animal Plant Health Agency. Pigs were screened for absence of influenza infection by hemagglutination inhibition using four swine IAV antigens. Pigs were randomly divided into two groups and were inoculated intranasally with 2.2 x 10^5^ PFU virus. Previous experiments showed that the standard deviation in viral shedding within groups is around 0.8 log_10_ pfu/ml, so a group size of five pigs was sufficient to detect a difference in viral shedding with 80% power and 95% confidence (power and sample size calculation for a one-way ANOVA with three groups in Minitab 17). Two milliliters were administered to each nostril using a MAD300 mucosal atomization device (Wolfe Tory Medical). Four nasal swabs (two per nostril) were taken daily after the challenge. Two nasal swabs were placed in 2 ml virus transport medium for the quantification of viral load by plaque assay as previously described (63). The other two nasal swab samples were put directly into TRIzol (Invitrogen, ThermoFisher Scientific, UK) for subsequent RNA isolation according to the manufacturer’s instructions. Animals were humanely killed 5 d post challenge. At post mortem, the lungs were removed, and photographs taken of the dorsal and ventral aspects. Macroscopic pathology was scored blind, as previously reported (64). Tracheal swabs, bronchoalveolar lavage and the accessory lobe were collected to determine the viral load in the lower respiratory tract by plaque assay. In brief, the accessory lobe was cut into small pieces and homogenised with a gentleMACS Octo Dissociator (Miltenyi Biotec) using C tubes (Miltenyi Biotec) in ice cold Dulbecco’s PBS supplemented with 0.1% BSA. The lung homogenates (10% w/v) were centrifuged and the clarified supernatant was used to determine the viral load and for RNA isolation. The tracheal swabs were processed like the nasal swabs. BALF was collected as previously described (63) and cell-free supernatant used to determine viral load. For histopathology, five lung tissue samples per animal from the right lung (two pieces from the apical, one from the medial, one from the diaphragmatic and one from the accessory lobe) were collected into 10% neutral buffered formalin for routine histological processing at the University of Surrey. Formalin-fixed tissues were paraffin wax–embedded, and 4 µm sections were cut and stained with H&E. Immunohistochemical staining of IAV NP was performed in 4 µm tissue sections as previously described (65). Histopathological changes in the stained lung tissue sections were scored by a veterinary pathologist blinded to the treatment group. Lung histopathology was scored using five parameters (necrosis of the bronchiolar epithelium, airway inflammation, perivascular/bronchiolar cuffing, alveolar exudates and septal inflammation) scored on a five-point scale of 0 to 4 and then summed to give a total slide score ranging from 0 to 20 and a total animal score from 0 to 100 (66). The Iowa system includes both histological lesions and immunohistochemical staining for NP (35). Sequencing of segment 5 from virus-positive samples from pigs confirmed that viruses produced during infection encoded uAUG or not, as expected (data not shown).

### Numerical analyses

The numbers of replicates for each experiment are defined in the figure legends. Independent experiments are defined as replicates carried out on different days. Scatter plot data points indicate data points from individual samples. Statistical analyses were chosen according to published advice (67) and unless otherwise stated, were performed using Graphpad Prism.

## Acknowledgements

We thank Dr Ben Killingley for clinical virus isolates, Dr Ed Hutchinson for critical comments, the Easter Bush Pathology Service staff for assistance and the animal staff at the Roslin and Pirbright Institutes and University of Ottawa for animal care.

## Supplementary Information

**Table S1.** Distribution of NP start codons by major IAV lineage in the stratified subsampled data set.

| Host-lineage | Avian | Swine | Human | Other | Total Sequences | Conventional Start % | uAUG Start % |
| --- | --- | --- | --- | --- | --- | --- | --- |
| Avian-Americas | 2642 | 2 | 2 | 5 | 2651 | 99.9 | 0.1 |
| Avian-Eurasia | 1449 | 22 | 56 | 46 | 1573 | 99.6 | 0.4 |
| Equine-Canine | 0 | 2 | 0 | 71 | 73 | 100.0 | 0.0 |
| Human-pdm2009 | 2 | 6 | 232 | 0 | 240 | 2.1 | 97.9 |
| Human-Seasonal | 1 | 28 | 894 | 1 | 924 | 99.6 | 0.4 |
| Swine-Classical-Triple | 15 | 352 | 23 | 0 | 390 | 8.7 | 91.3 |
| Swine-Eurasia | 1 | 68 | 4 | 0 | 73 | 94.5 | 5.5 |
| Variant-pdm2009 | 2 | 122 | 181 | 4 | 309 | 7.8 | 92.2 |
| Unclassified | 2 | 1 | 0 | 6 | 9 | 100.0 | 0.0 |

**Table S2. (see separate Excel file).** Stratified subsampled dataset of IAV segment 5 sequences, classified according to the presence (TRUE) or absence (FALSE) of an inframe upstream AUG codon in the NP gene, as well as H and N subtype, continent, region and country of isolation, host and date of isolation and named clade. The position of the first AUG codon in the segment (first_start) and the identity of the nucleotide triplet at the location of the uAUG (upstream3) are also tabulated.

**Figure S1.**
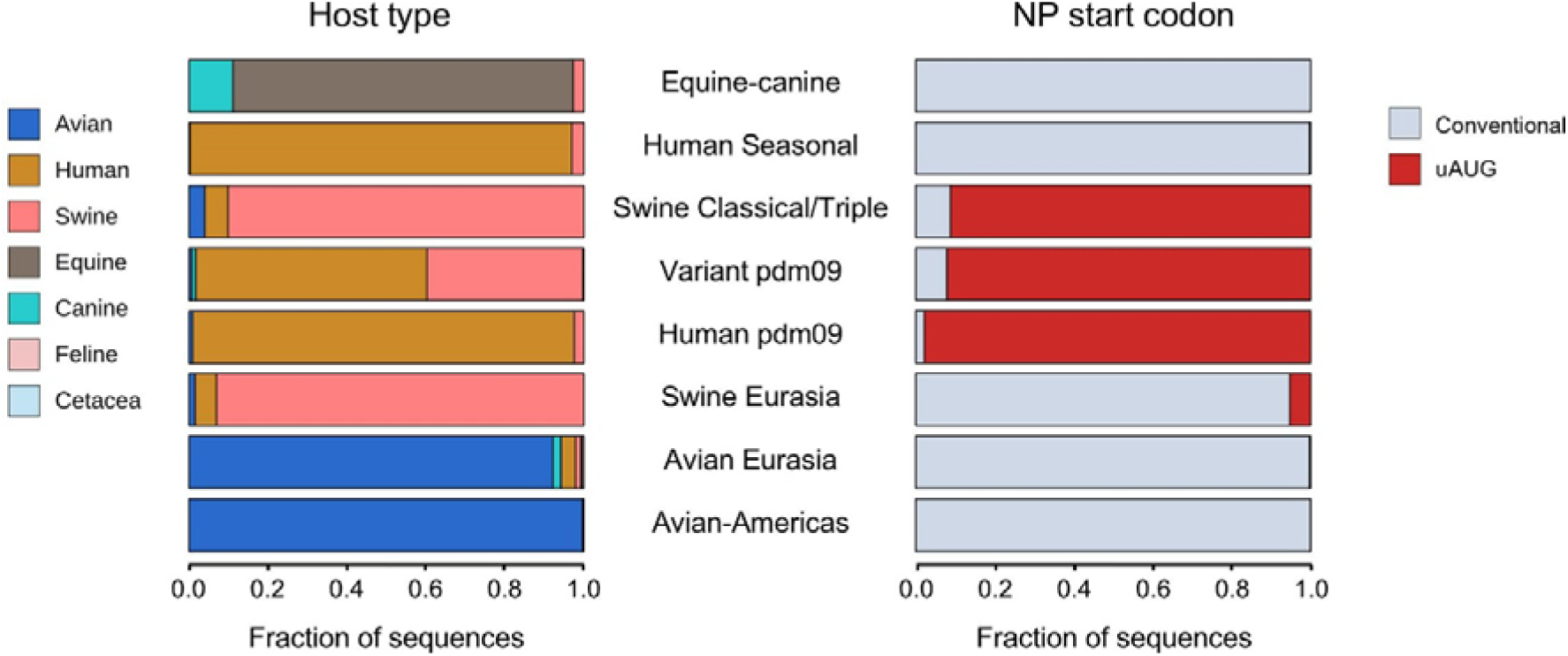
Distribution of hosts and segment 5 start codon positions by major IAV lineage in the stratified subsampled data set.

**Figure S2.**
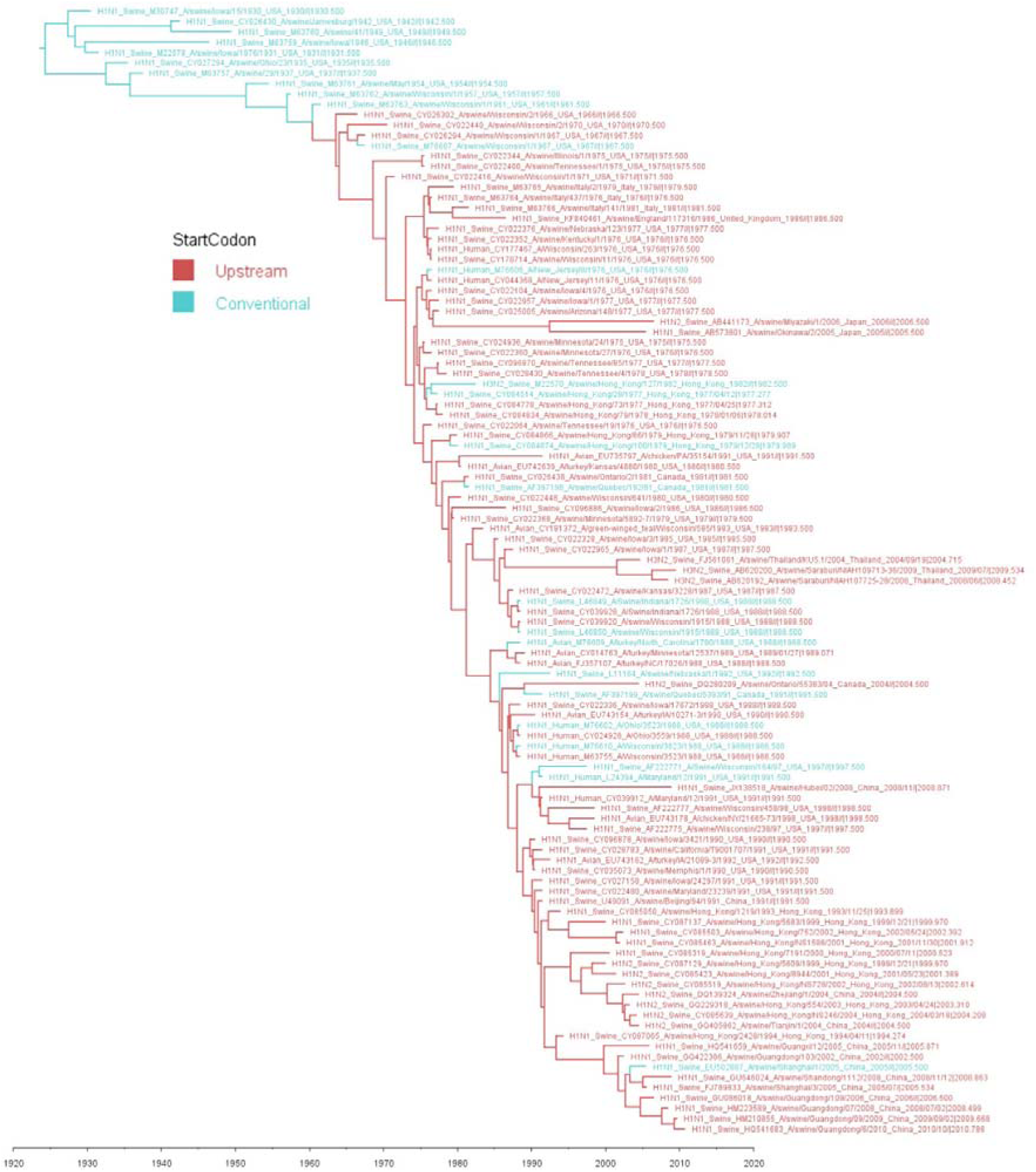
Time scaled tree of segment 5 from the classical swine virus clade with skygrid tree prior and NP start codon as a discrete trait. Sequences in red possess the uAUG, those in blue do not. The uAUG codon first appeared at an estimated date in 1962 (range 1958-1965) in H1N1 swine IAV in Wisconsin. This is based on the mutation happening somewhere on the branch between nodes 1960.41 [Highest Posterior Density (HPD): 1958.1 - 1961.5] and 1963.55 [HPD: 1961.2 - 1965.7]).

**Figure S3.**
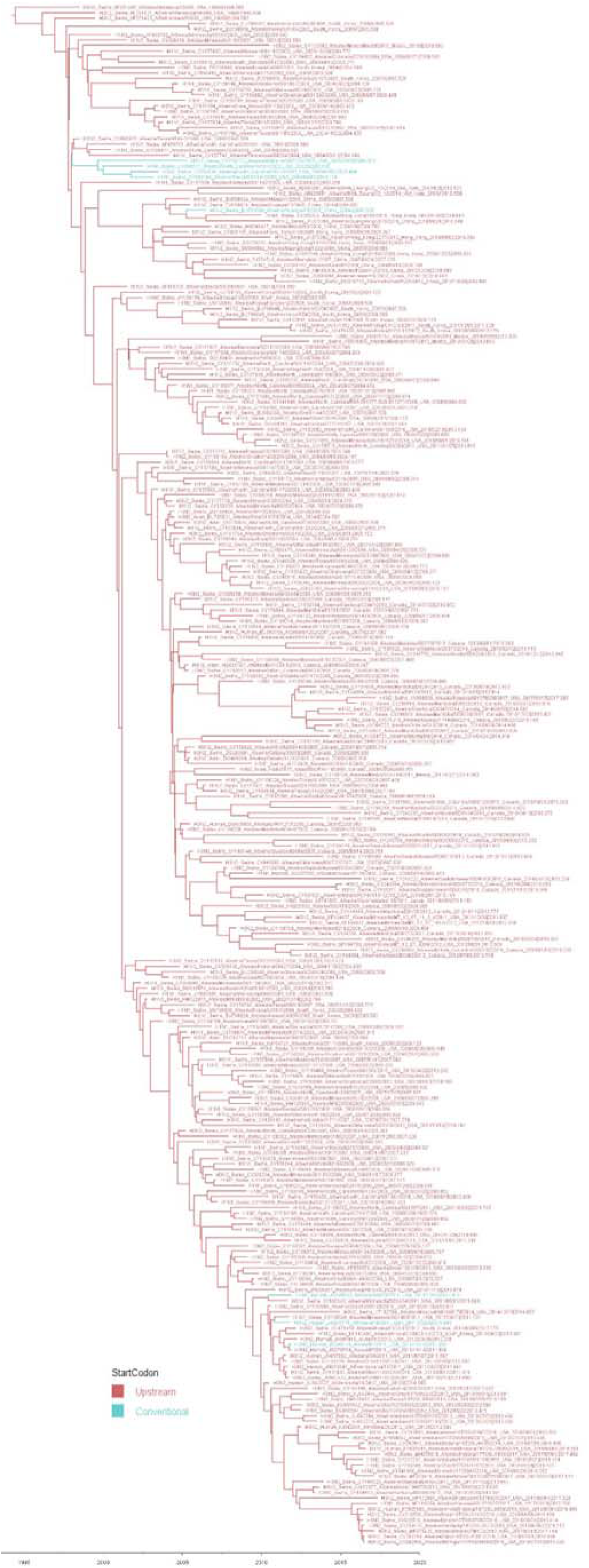
Time scaled tree of segment 5 from the triple reassortant swine virus clade with skygrid tree prior and NP start codon as a discrete trait. Sequences in red possess the uAUG, those in blue do not. Triple reassortant viruses most likely inherited an uAUG-containing segment 5 from their classical swine progenitor and subsequently, have largely kept it.

**Table S3.** Sporadic occurrences of the uAUG in avian and human seasonal viruses. Isolate name, subtype, segment 5 sequence accession code, host and country of isolation are tabulated. Most isolates are too temporally and/or geographically separate to represent linked events; the H6 duck isolates from the eastern USA in 1999 and 2002 and the human H3N2 isolates from the mid 1970s might be exceptions.

| Strain | Accession | Host | Subtype | Country |
| --- | --- | --- | --- | --- |
| A/duck/New_York/16873/1999 | CY014891 | Avian | H6N2 | USA |
| A/mallard/Maryland/887/2002 | EU026010 | Avian | H6N1 | USA |
| A/chicken/Italy/322/2001 | CY021552 | Avian | H7N1 | Italy |
| A/chicken/Israel/702/2008 | GQ148843 | Avian | H9N2 | Israel |
| A/duck/Taiwan/DC167/2010 | KC693624 | Avian | H1N3 | Taiwan |
| A/Albany/20/1974 | CY021096 | Human | H3N2 | USA |
| A/Bilthoven/2271/1976 | KC296468 | Human | H3N2 | Netherlands |
| A/Singapore/64K/2007 | KP223188 | Human | H3N2 | Singapore |
| A/Victoria/600/2016 | CY254980 | Human | H3N2 | Australia |

**Figure S4.**
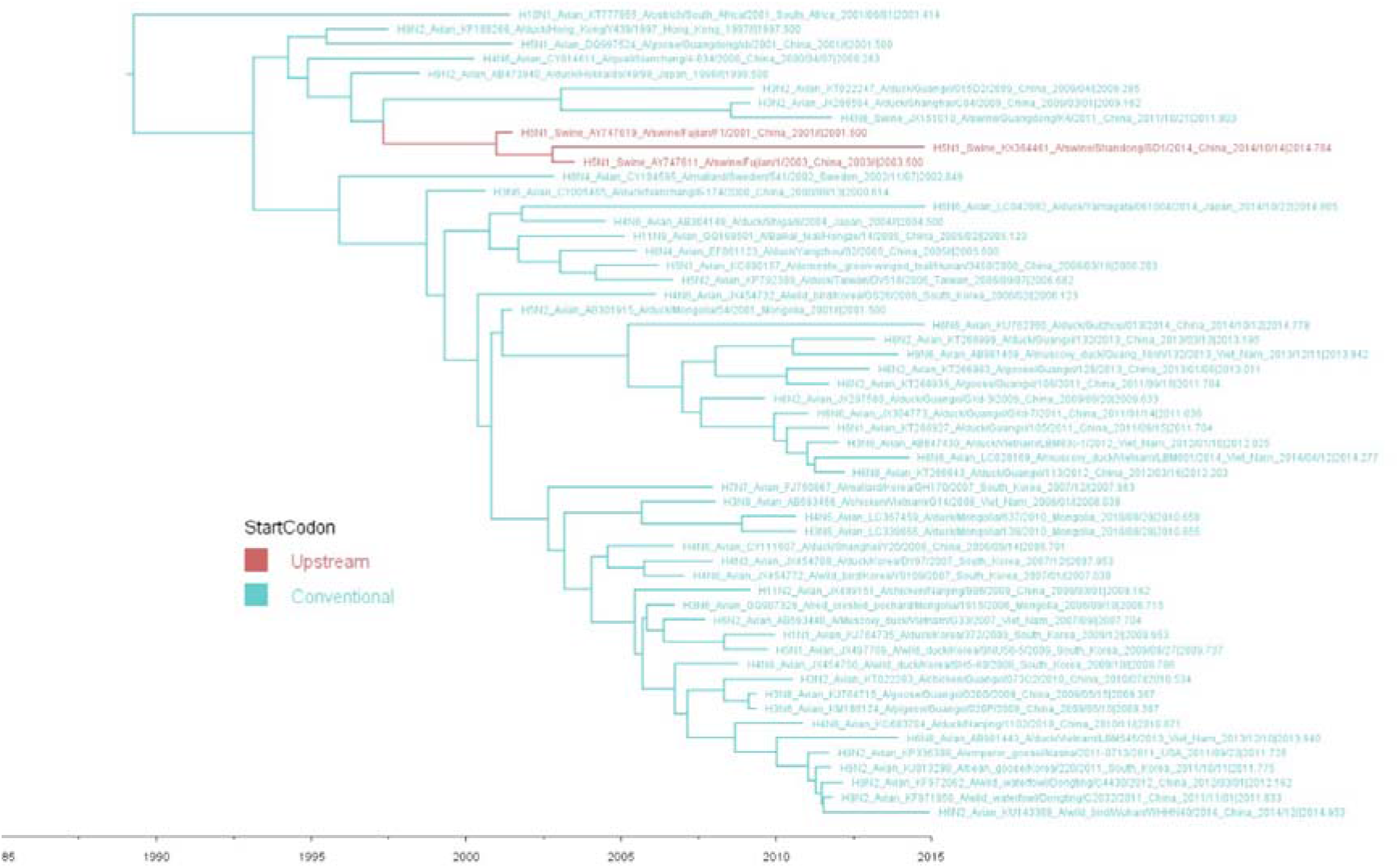
Acquisition of the segment 5 uAUG codon associated with epizootic transfer from ducks to swine. A time scaled tree of segment 5 from an avian virus clade with skygrid tree prior and NP start codon as a discrete trait. Sequences in red possess the uAUG, those in blue do not. The 5’-UTR sequence of segment 5 from the closest relative of the uAUG-possessing A/swine/Fujian/2001 and /2003 viruses (A/duck/Zhejiang/11/2000(H5N1)) has not been reported, making it uncertain whether the polymorphism occurred before or after the host-range jump. Note also that the apparent persistence of the uAUG-containing swine virus in China until 2014 may be an artefact, as all eight segments of A/swine/Shandong/SD1/2014 have the corresponding genes from A/swine/Fujian viruses from the early 2000s as their closest relatives (> 99.7% nucleotide identity) and conversely, lack close relatives from the 2010s, raising the possibility of laboratory contamination.

**Figure S5.**
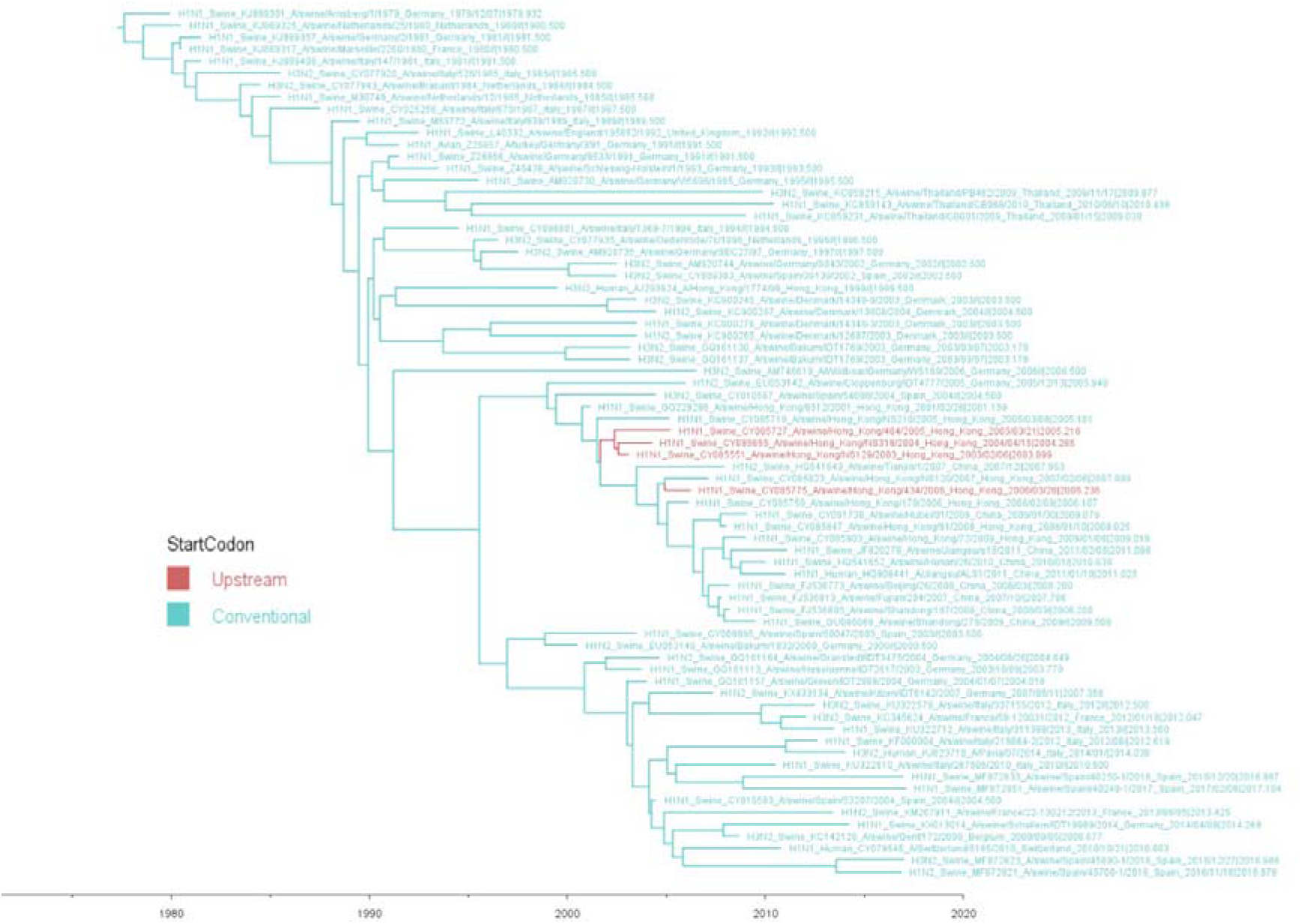
Acquisition of the segment 5 uAUG codon within the Eurasian swine IAV lineage. A time scaled tree of segment 5 from an avian virus clade with skygrid tree prior and NP start codon as a discrete trait. Sequences in red possess the uAUG, those in blue do not.

**Table S4.** Summary statistics for minireplicon assay curve fitting data reported in Figures 3C, D. See separate Excel file.

**Figure S6.**
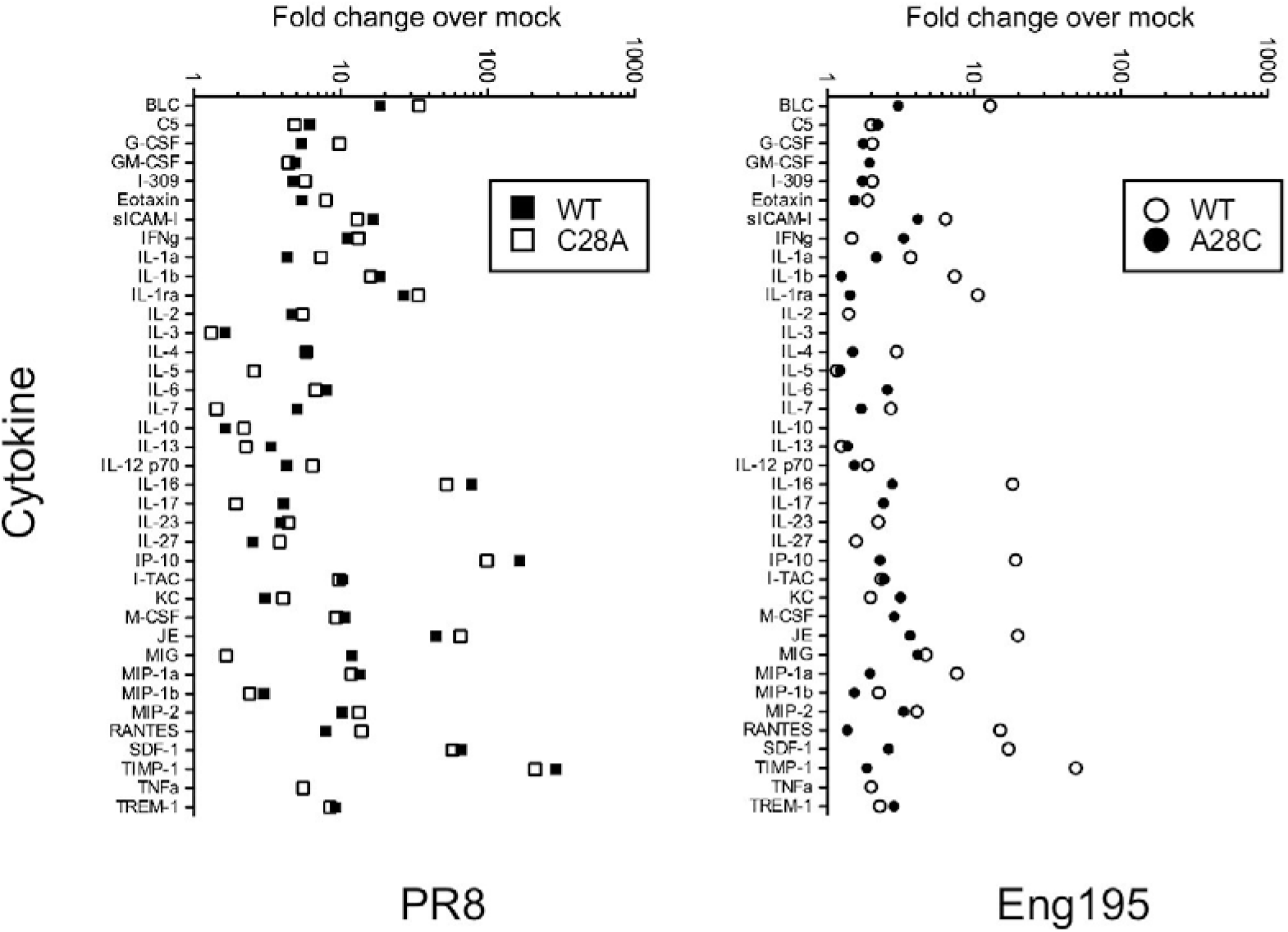
Cytokine levels in infected mouse lung. Lung homogenates from mice infected with the indicated viruses or mock infected animals were pooled and the levels of various cytokines measured by cytokine array.

**Table S5. (see separate Excel file).** Formalin-fixed lung sections were stained with haematoxylin and eosin and examined by a veterinary pathologist. Six pathological changes (epithelial cell degeneration and necrosis, perivascular inflammation, peribronchial inflammation, interstitial inflammation, interstitial necrosis and lymphocyte cuffing) were scored on a scale of 0-4. The percentage of lung affected was estimated visually. The pathological changes present along with a consideration of the area of lung affected was used to give an overall qualitative score of the severity of histopathological changes. Slides from three mice (22, 14 and 9) were not scored and excluded from the analysis due to marked atelectasis (artefact) which precluded assessment of pathological changes.

**Figure S7.**
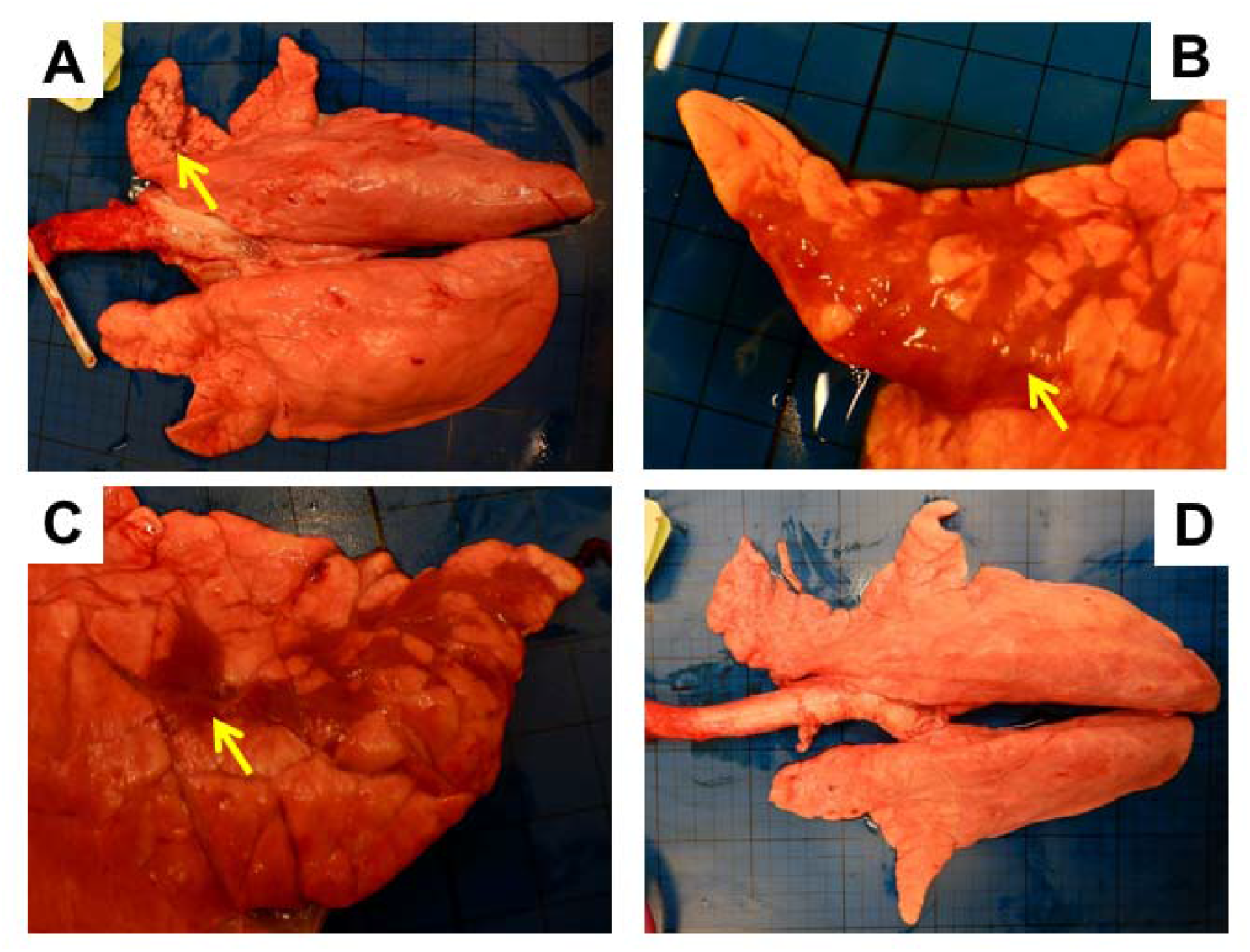
Macroscopic pathology of pig lungs 4 days post challenge with WT (A, B and C) and A28C Eng195 (D) viruses. Arrows indicate areas of atelectasis observed in the apical lobe of the right lung (A) and medial lobes (B and C). Some animals from the A28C group exhibited no remarkable gross pathology (D).

**Table S6. (see separate Excel file)**. Histopathological changes in the stained lung tissue sections were scored by a veterinary pathologist blinded to the treatment group. Lung histopathology was scored using five parameters (necrosis of the bronchiolar epithelium, airway inflammation, perivascular/bronchiolar cuffing, alveolar exudates and septal inflammation) scored on a five-point scale of 0 to 4 and then summed to give a total slide score ranging from 0 to 20 and a total animal score from 0 to 100 (63). The Iowa system includes both histological lesions and immunohistochemical staining for NP (35).

